# Astrocytic glutamate regulation is shaped by adversity and glucocorticoid signalling

**DOI:** 10.1101/2024.06.24.600362

**Authors:** Dominic Kaul, Amber R Curry, Nathalie Gerstner, Anna S Fröhlich, Anthi C Krontira, Christina Kyrousi, Caine C Smith, Greg T Sutherland, Mirella Dottori, Michael J Ziller, Christos Chatzinakos, Nikolaos P Daskalakis, Silvia Cappello, Darina Czamara, Elisabeth B Binder, Cristiana Cruceanu, Janine Knauer-Arloth, Naguib Mechawar, Sibylle G Schwab, Lezanne Ooi, Natalie Matosin

## Abstract

Astrocytes are a brain cell type vulnerable to the effects of stress and the development of psychiatric-like phenotypes in animals, yet how this translates to humans is unclear. Here, we probed the diversity of ∼145,000 total human cortical astrocytes with single nucleus and spatial transcriptomics, showing that human astrocytes comprise a molecularly and anatomically diverse cell population. In individuals with psychiatric disorders and high adversity exposure, we identified distinct alterations to glutamate-related synaptic functions, supported by histological quantification of >20,000 astrocytes. Early-life adversity exposure produced more pronounced cellular changes than adversity experienced later in life, and female cases displayed stronger transcriptomic associations than males with adversity exposure. Human pluripotent stem cell-derived astrocytes from both two- and three-dimensional models confirmed that glutamate signalling is directly impacted by glucocorticoid activation. Our findings highlight astrocytes as crucial players in how exposure to severe adversity raises risk to psychopathology and position them as strategic pharmacological targets for future intervention strategies.

---

The accumulation of recent geopolitical, environmental, and health crises has highlighted that exposure to severe adversity leading to prolonged biological stress has broad and substantial ramifications for global mental health. For example, in the first year of the COVID-19 pandemic, the worldwide incidence of depression and anxiety increased by 25%^1^. It is also estimated that at least one in two children is impacted by maltreatment, a major psychological stressor, globally^2^. Such adversity exposures are robust risk factors for a plethora of medical conditions, including psychiatric disorders^3,4^, and are often associated with impaired treatment responsiveness^5,6^ and increased symptom severity^7-9^. Despite strong evidence that stress plays a clinically significant biological role in mental health disorders, identifying its molecular effects in human populations remains challenging due to difficulty defining “impactful” adversity exposure and linking it reliably to specific clinical outcomes, such as diagnosis. In both rodents and non-human primates, recent approaches have begun to uncover molecular signatures of adversity and highlight that both cell-type^10,11^ and spatial domains^12^ of the prefrontal cortex (PFC) are essential to shaping the brain in response to biological stress. Astrocytes are one such cell type that have emerged as a recent focus in psychiatric disorders^10,13,14^. Cerebral astrocytes are highly abundant and play essential roles across many brain functions, especially in coordinating the synaptic, metabolic, and neuroinflammatory environments^15^. Astrocytes also directly contact blood vessels, positioning them as key responders to circulating signalling molecules passing the blood-brain barrier, including cortisol^16^. However, astrocytes represent a molecularly and regionally diverse population of cells^16-19^ and how adversity exposure shapes this diversity in the human brain, particularly in the context of psychiatric disorders, is unknown.

Here, we performed comprehensive profiling of human cortical astrocytes with a range of high-resolution sequencing techniques, including single nucleus RNA sequencing (snRNAseq) of ∼800,000 cells and over 110,000 astrocytes from the postmortem orbitofrontal cortex (OFC) of 87 individuals. Spatial transcriptomics was also performed using a subset of 13 of these individuals^20,21^. We also analysed the morphology of more than 20,000 astrocytes to explore the molecular and spatial diversity of astrocytes in this neocortical area. We focus on the OFC as it is important to emotion-related learning and perceiving adversity^22^. It is also implicated in the pathology of psychiatric disorders^23,24^ offering unique insight at the intersection of biological stress and psychiatric diagnosis. We first determined the molecular and spatial profile of astrocyte populations in the OFC. We then examined this cohort to identify how cases of psychopathologies with high adversity exposure before diagnosis vary from those without, and replicate these findings in a second cohort. We focused on dissecting the persisting alterations to astrocytes associated with significant adversity exposure and explored the importance of sex and timing of adversity exposure (childhood vs adulthood). We also demonstrated that core functions of astrocytes that regulate glutamate neurotransmission are responsive to the activation of cortisol receptors by employing multiple human pluripotent stem cell (hPSC)-derived astrocyte models. Together, transcriptomic, morphological, and *in vitro* evidence converge to show that human astrocytes consistently change due to biological stress triggered by exposure to adversity. This has the potential to guide treatment specificity as well as the development and implementation of early intervention and prevention strategies for psychiatric disorders related to severe adversity exposure.

## Results

### SnRNAseq identifies transcriptional diversity of astrocytes in the human orbitofrontal cortex

To investigate the impact of adversity on the population of human astrocytes, we analysed postmortem OFC (Brodmann Area 11) samples derived from 32 neurotypical individuals and 55 individuals diagnosed with a psychiatric disorder (major depression, bipolar disorder, schizoaffective disorder, or schizophrenia; Figure 1; demographics outlined in Supplementary Figure 1). We have previously sequenced 787,046 nuclei from these samples,^25^ with ∼13% of nuclei (111,516) labelled as astrocytes (Figure 2a). Astrocytes were also previously clustered at low-resolution as either ‘protoplasmic’ (grey matter; *ATP1A2, GJA1* and *SGCD*), or ‘fibrous’ (white matter; *GFAP, ARHGEF4*)^25^. To first explore astrocyte diversity at high-resolution, we performed Leiden clustering restricted to nuclei labelled as astrocytes. This identified 10 astrocyte sub-clusters (Figure 2a, Supplementary Figure 2d-f). Several aligned with astrocyte sub-clusters identified elsewhere. This included a cluster enriched for amyloid metabolism (*APOE* and *CLU*; cluster 1), ‘pan-reactive’-like astrocytes (*GFAP, VIM, NEAT1*; cluster 8), and a small population of phagocytotic astrocytes (*ELMO1, MEF2A*; cluster 9) seen in an integration of 17 human snRNAseq datasets across 302 individuals^26^. We also identified clusters demarcated by *LSAMP* and *GPC5* (cluster 3) or *MALAT1* and *DPP10* (cluster 6) that were seen in a large hippocampal dataset^27^ (Figure 2b). We also identified cortex-specific interlaminar-like astrocytes, enriched for the markers *ID3, SERPINI2*, and *WDR49* (cluster 6; Supplementary Data 1)^13,28^. Astrocyte sub-clusters were largely divided between grey matter ‘protoplasmic’ clusters 0, 1, 2, 3, and 5 (88% of the protoplasmic label) and white matter ‘fibrous’ clusters 4, 6, and 7 (98% of the fibrous label). The exception was reactive-like cluster 8 being identified in both low-resolution annotations (84% protoplasmic, 16% fibrous; Supplementary Figure 3). This supports that general localisation to grey/white matter is the strongest driver of molecular differences in astrocytes, but that further diversity exists within these domains.

**Figure 1:**
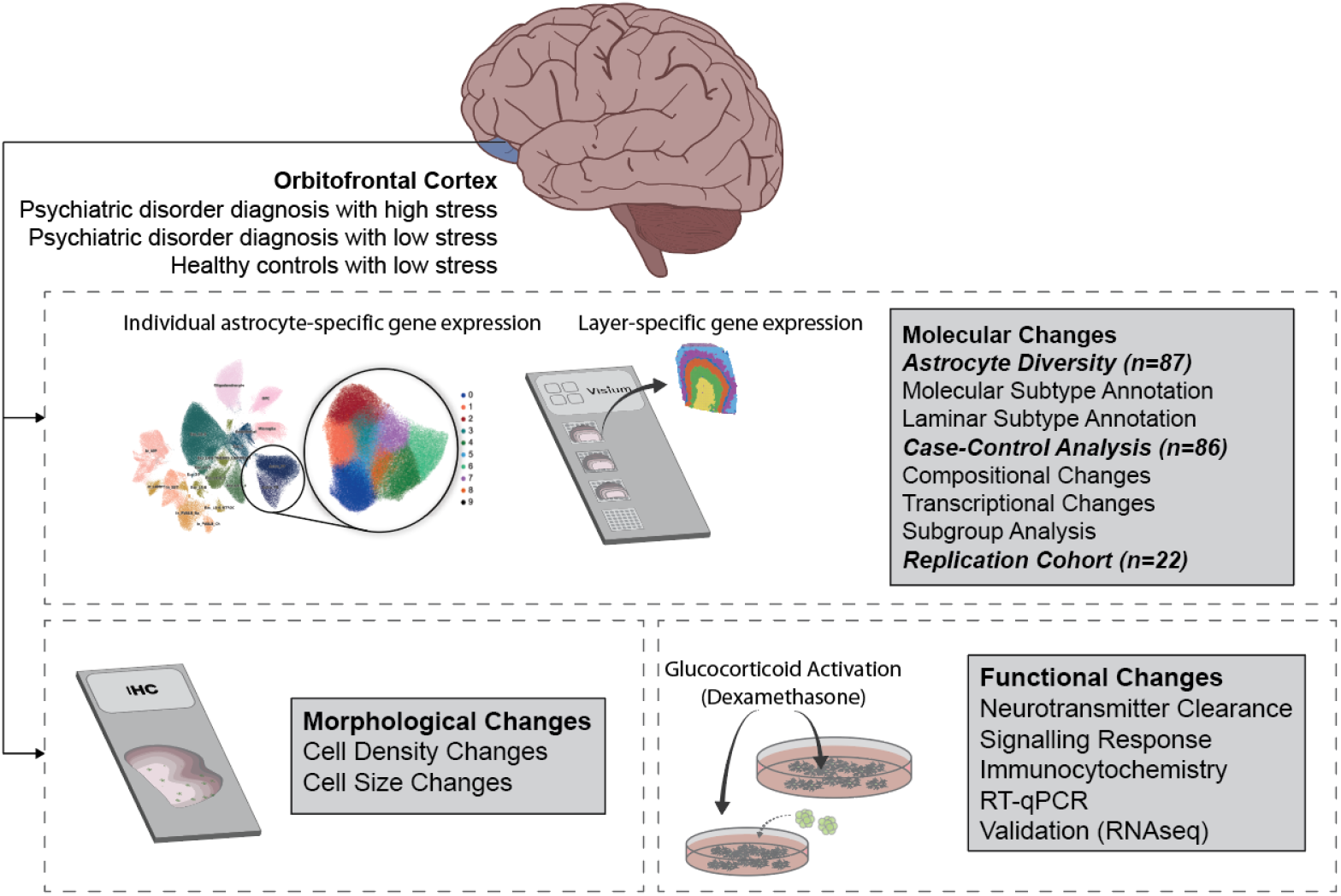
Schematic of experimental workflow. Profound history of stress and previous diagnosis of severe psychiatric disorder were extracted from a postmortem cohort of 92 individuals. Tissue sections from the 3rd ∼1cm coronal slice of Brodmann area 11 were acquired. 787,046 nuclei (111,516 astrocytes) were extracted, processed, sequenced, integrated, and then labelled by cell type, consisting of 87 libraries which met quality control thresholds. Additional sections were mounted onto Visium (10x Genomics) slides for spatial transcriptomics from a subset of the 87 individuals with libraries (n=13). A replication cohort of 22 individuals was also used to validate findings. Immunohistochemistry in a largely overlapping subset of this cohort was used to validate spatially annotated astrocyte populations associated with a history of profound stress. Glucocorticoid activation of human pluripotent stem cell-derived astrocytes was used to determine alterations to the synaptic functions.

**Figure 2:**
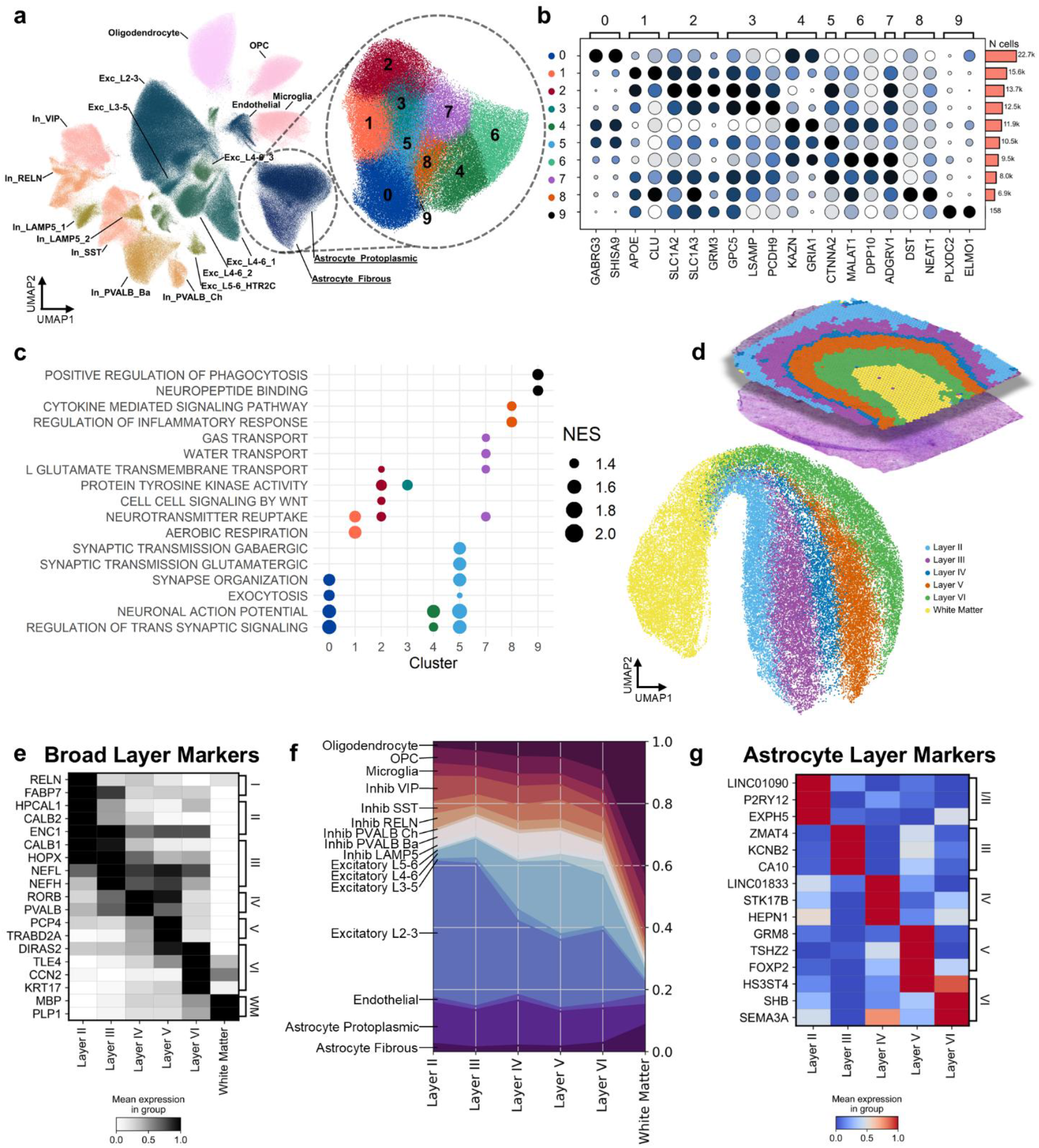
Characterisation of human astrocytes in the orbitofrontal cortex highlights functional and regional heterogeneity. **(a)** UMAP representation of single nucleus RNA sequencing (snRNAseq) from 787,046 nuclei and 111,516 reclustered astrocytes (in zoom) coloured by cluster as determined by Leiden algorithm. Fibrous astrocyte markers: *GFAP, ARHGEF4*, Protoplasmic astrocyte markers: *ATP1A2, GJA1, SGCD* **(b)** Dotplot of marker genes for each astrocyte cluster from snRNAseq analysis. Dot colour indicates scaled mean gene expression across clusters and dot size indicates proportion of nuclei in cluster above single cell transformed expression (cutoff = 0.5). **(c)** Gene set enrichment analysis using the Biological Processes database by cluster. Ranks were the score derived from the scanpy function rank_gene_markers, method = Wilcoxon. Terms were pruned for semantic similarity using the GO.db to keep only the most significant child terms. Dots indicate significant enrichment per cluster (P_adj_ < 0.05), size indicates normalised enrichment score, and colour indicates cluster. **(d)** UMAP representation of spatial transcriptomics (Visium) dataset from 13 individuals consisting of ∼50,000 spatial points. Integration and clustering between individuals were performed using STAligner and mclust pipelines (upper). Datapoints from integrated dataset were mapped onto individual slides to confirm regional identities (lower). **(e)** Broad cortical layer markers from spatial transcriptomics curated from manually annotated datasets. Darkness indicates mean expression in group, scaled per gene. **(f)** Integration of the complete snRNAseq dataset with the complete spatial transcriptomics dataset using Tangram. Colours represent the proportion of each cell type at each layer of the cortex. **(g)** Layer markers for astrocytes. Cells were defined into layers using the Tangram tool (>1.5-fold change confidence), and pseudobulk counts per individual were used to determine candidate layer markers. Colour indicates mean expression in group, scaled per gene.

Next, we performed gene set enrichment analysis (GSEA) across the transcriptome of each sub-cluster to explore whether transcriptionally defined populations of astrocytes associated with distinct functional niches. Several sub-clusters associated with distinct astrocyte functions, including regulating synaptic activity (clusters 0 and 5; e.g. synapse organisation and exocytosis, reflective of clusters identified previously^18^), neurotransmitter and metabolic homeostasis (clusters 1, and 2 ; e.g. neurotransmitter reuptake) or immune response (cluster 8 and 9; e.g. cytokine signalling, and phagocytosis) (Figure 2c). Cluster 5 was enriched for glutamate and γ-aminobutyric acid (GABA) signalling, and cluster 1 was enriched for aerobic respiration. Further, cluster 7, which was enriched for both *SLC1A2* and *GFAP*, was uniquely enriched for functions related to water and gas transport. This cluster also expressed the water channel *AQP4*, and may represent varicose projection astrocytes, a substate of deeper layer/white matter astrocytes^29^. Across all clusters, the enriched functional terms were disproportionately associated with regulating the neuronal environment and neurotransmitter transport (59% of terms), supporting prior evidence that the surrounding cellular and synaptic environment is an important driver of astrocyte diversity (Supplementary Figure 3c-d, Supplementary Data 2)^30,31^.

### Spatial transcriptomics reveals that OFC astrocytes are anatomically diverse

Given previous evidence that astrocytes demonstrate functionally relevant laminar distributions in the cortex^30-32^, we also sought to determine whether we could detect laminar populations of astrocytes. To synthesise a transcriptional reference for these layers, we performed spatial transcriptomics (Visium, 10x Genomics) in a subset of individuals from this cohort (n=13, full demographics available in Supplementary Figure 1). After quality control, integration, and clustering, we delineated six spatial domains across all 13 individuals (layers II, III, IV, V, VI, and white matter; Figure 2d). Clustering did not associate with covariates (Age, Sex, PMI, Batch), and domains were consistent between individuals when domains were remapped onto individual slides *in situ* (Supplementary Figure 4). Layer markers were then corroborated with prior spatial transcriptomics datasets that manually annotated layers based on histology (layer I: *RELN*, layer II: *CALB1*, layer III: *NEFL*, layer IV: *RORB*, layer V: *PCP4*, layer 5: *DIRAS2*, white matter: *MBP*; Figure 2e, top 100 genes per layer are provided in Supplementary Data S3)^33,34^. We also performed gene ontology analysis on the top 100 genes from each layer using EnrichR, which further supported layer identities, highlighting domains enriched for relevant functions (e.g. white matter: myelination, layer V: signal release from synapse, layer III: GABAergic synaptic transmission, Supplementary Data S4). Together, this semi-automated integration method identified conserved histologically relevant domains across the human OFC.

Previous spatial mapping of broad astrocyte clusters (e.g. fibrous vs protoplasmic) using snRNAseq deconvolution has largely assigned astrocytes to either layer I or white matter^35,36^, likely given their proportional abundance in these regions. To more finely classify astrocytes to their layer of origin, we employed the cluster mode of Tangram to estimate the probability of each cell occurring at each annotated layer defined from all 13 individuals. We only mapped an astrocyte if it was 1.5 times likely to appear one layer, relative to all other layers. Using this pool of astrocytes, we identified several candidate molecular markers for well-defined laminar astrocyte populations, including semaphorin 3A (*SEMA3A*) in layer VI and a potassium voltage-gated channel subunit (*KCNB2*) in layer III (Figure 2g; Supplementary Data 5). Across the layers of grey matter, most sub-clusters were not biased towards a specific layer, except cluster 2, involved in neurotransmitter homeostatic functions, which was enriched in layer IV a primary input layer of the cortex (36.2% of astrocytes in this layer). The white matter was appropriately represented by clusters 4 and 6 (65% of astrocytes in this region). However, it is important to note that only 56% of astrocytes met the threshold applied, suggesting that transcriptionally distinct laminar populations identified remain modest with this method. Together, these approaches demonstrate that astrocytes are transcriptionally diverse in the human cortex and that cell state and spatial localisation influence this diversity.

### High adversity exposure delineates cluster-specific transcriptomic changes in grey matter astrocytes

Exposure to adversity is a trans-diagnostic environmental risk factor for psychiatric disorders. However, high-resolution profiling of molecular markers that underlie this shared risk remain unknown. Using this foundation of astrocyte diversity, we aimed to explore the trans-diagnostic impact of adversity exposure on astrocytes. Individuals diagnosed with psychiatric disorders were stratified to either those exposed to high adversity before diagnosis (three bipolar disorder/four major depression/two schizoaffective disorder/15 schizophrenia) vs low adversity (two bipolar disorder/two major depression/five schizoaffective disorder/23 schizophrenia), based on clinical history. For simplicity, we herein refer to these high adversity and low adversity, respectively. Unfortunately, we did not have access to high adversity controls. To validate effects associated with psychiatric diagnosis, both groups were also contrasted against the group of case and control individuals with low adversity. This resulted in three groups: high-adversity-cases (n=24), low-adversity-cases (n=32), low-adversity-controls (n=31).

Using the spatially labelled cells, there were only minor differences in the composition of astrocytes assigned to each layer in the high-adversity-cases (Figure 3c) compared to low-adversity-cases (Figure 3b) and low-adversity-controls (Figure 3a). Both case groups had an increase in the proportion of cluster 2 increased by 6.6-7.0% in layer IV, compared to controls. Cluster 1 was also notably decreased (9.1%) in layer II in high-adversity-cases compared to low-adversity-cases. In support of this effect, in the snRNAseq dataset, there was a nominal decrease in the proportion of protoplasmic astrocytes across broad cell types (P=0.023), as well as a nominally significant increase within astrocyte cluster 2 (P=0.012) between low-adversity-cases and low-adversity-controls and a decrease between the high-adversity-cases and low-adversity-cases in cluster 1 (P=0.045), although these did not survive correcting for multiple testing (Padj>0.05; Supplementary Figure 5).

**Figure 3:**
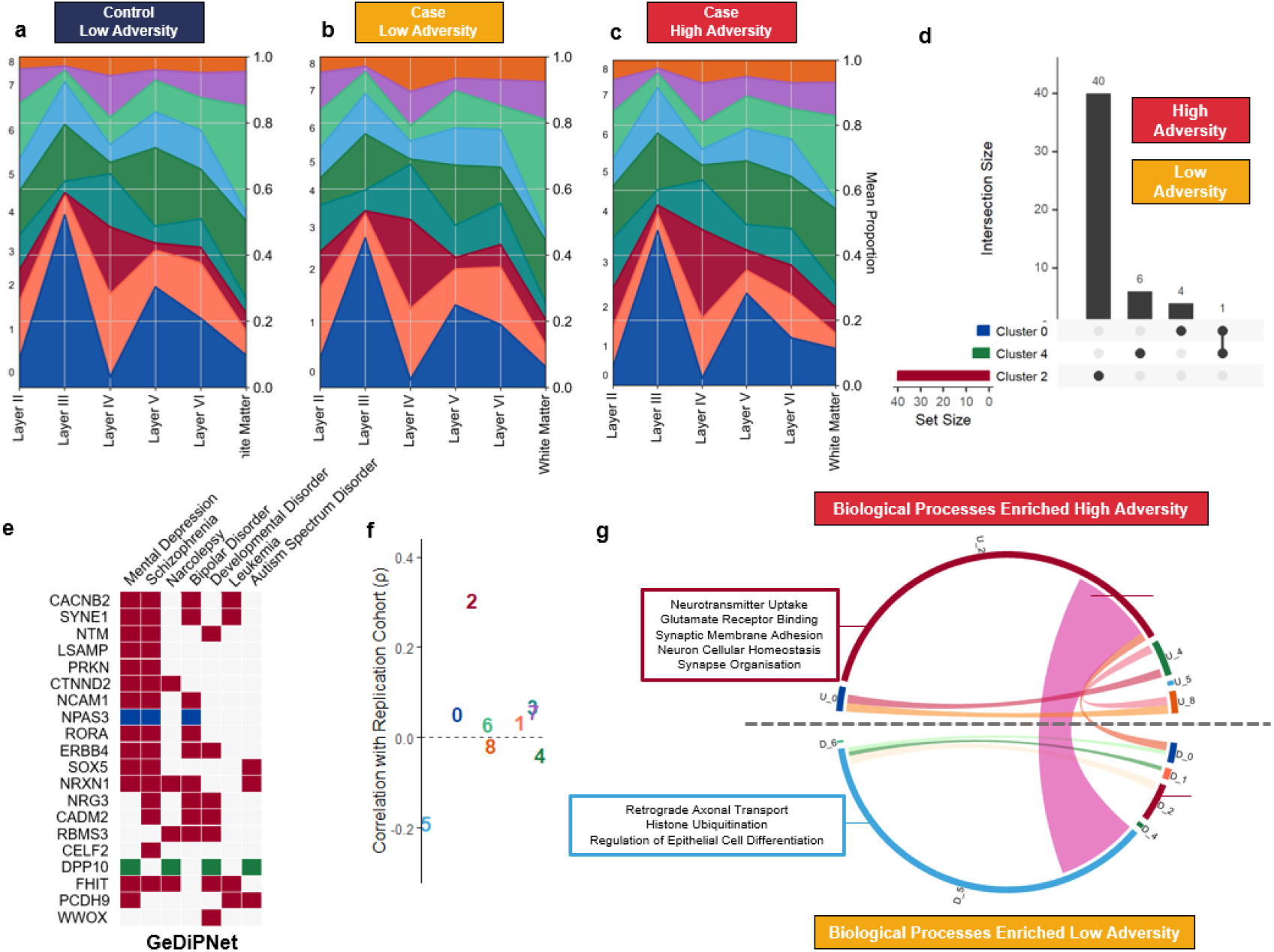
Cortical astrocyte diversity delineates cases of psychiatric disorders with a history of high adversity exposure. (a-c) Integration of snRNAseq astrocyte-specific dataset from adversity delineated groups with spatial transcriptomics. Datasets were integrated using the Tangram clusters mode to predict cell likelihood at each laminar distribution. Graphs represent average proportion of cells per individual (i.e. each individual is weighted equally) (d) UpSet plot depicting the differentially expressed genes between psychiatric disorder cases stratified by history of adversity exposure (high vs low; Padj<0.1). Set size indicates number of genes meeting threshold for each cluster and dots connected by a line indicate common genes. (e) Heatmap of gene-disease association. DeDiPNet 2023 database, using the EnrichR tool. Colour indicates cluster of origin. (f) Correlation of gene ranks (sign(logFC) * -log10(P)) between clusters in primary OFC cohort and replication DLPFC PTSD cohort (Spearman’s correlation). Colour indicates cluster. (g) Chord diagram of gene set enrichment analysis performed on differentially expressed gene lists per cluster. Size of arc denotes number of significant terms (Padj<0.05) and ribbons denote common terms. GO Terms delineated based on positive (U_) or negative (D_) enrichment in cluster. Candidate functional terms annotated following pruning for semantic similarity.

To capture transcriptomic changes associated with high adversity exposure, we performed differential expression analysis with a model fitting all groups, including both case and control groups. Pairwise comparisons were then extracted to compare between the individuals diagnosed with a psychiatric disorder stratified by adversity exposure (high-adversity-cases vs. low-adversity-cases; Figure 3). The most differentially expressed genes (DEGs, PFDR < 0.1) were identified in cluster 2 (40 DEGs; 33 upregulated), including increased expression of the nuclear receptor RORA, previously associated with childhood abuse in cases of post-traumatic stress disorder39 and Limbic System Associated Membrane Protein (LSAMP), which is known to be increased in the synaptosome of the OFC in postmortem individuals with schizophrenia40 (Figure 3d, Supplementary Data 6). Only a single gene, the cation channel TRPM3, was implicated in more than one cluster (downregulated in clusters 0 and 4, Figure 3d). We also identified that DEGs between cases and controls without a psychiatric disorder were broadly consistent regardless of adversity exposure (low-adversity-controls vs. low-adversity-cases or high-adversity-cases). These effects were similarly most pronounced in the homeostatic clusters 2, 3, and 7 (Supplementary Data 7). To support whether DEGs associated with high adversity were relevant to psychiatric disorders, we used the EnrichR tool to assess gene-disease association. Using the GeDiPNet 2023 database, which contains over 7,000 diseases/disorders, mental depression (Padj=1.1×10-5, odds ratio=7.20) and schizophrenia (Padj=1.1×10-5, odds ratio=6.36) were the two most enriched disorders associated with stress-related DEGs (Figure 3e), supporting that the adversity exposure is relevant to these disorders.

To validate that the observed effects were driven by adversity, not specific diagnoses, we performed differential expression analysis in an independent snRNAseq dataset from a cohort consisting of postmortem dorsolateral PFC from individuals with post-traumatic stress disorder with high levels of complex childhood adversity (n=5) or adulthood adversity (n=6) (34,504 astrocytes), and low adversity exposed controls (n=11)10,41. Label transferring resulted in even allotment of the astrocytes across all clusters (0: 3588, 1: 5716, 2: 7463, 3: 3703, 4: 1872, 5: 979, 6: 2270, 7: 1754, 8: 6507). Comparing DEGs associated with prior high stress to the primary cohort using rank-rank hypergeometric overlap (RRHO), there was a moderate positive correlation in stress-related DEGs with cluster 2 from the primary cohort with cluster 2 from this second cohort (ρ = 0.30, P < 0.0001) but minimal correlation in all other clusters (Figure 3f). However, no DEGs survived FDR correction in the replication cohort, likely due its size.

We then performed GSEA to identify functions associated with high-adversity vs low-adversity cases within each cluster (Figure 3f), similar to recent analyses of psychiatric human postmortem cohorts10,14. This approach further spotlighted the functional enrichment changes in cluster 2, along with cluster 5 (146 and 125 biological processes Padj<0.05, respectively, Supplementary Data 8). Functions positively associated with high-adversity-cases in cluster 2 included increased neuron homeostatic functions and synaptic remodelling (synaptic membrane adhesion, and dendrite development) and specifically highlighted enrichment in the glutamate functions which demarcate this cluster (e.g. neurotransmitter uptake and glutamate receptor binding). In contrast, functions enriched in cluster 5 were associated with low-adversity-cases and included increased histone ubiquitination and cellular differentiation (Figure 3g). We also identified a module of functions enriched in diagnosis-high-stress in cluster 2 while also enriched in low-adversity-cases in cluster 5. This included functions related to WNT signalling, hormone stimulus, and cell morphogenesis. In the validation cohort, GSEA similarly identified functional enrichment for 65 out of 208 pathways that overlapped with those from the primary cohort (Supplementary Data 9). Of these, 26 were associated with cluster 2 and 23 with cluster 5. In cluster 2, the shared pathways included glutamate-related functions (such as glutamate receptor signalling, ligand-gated ion channel signalling, and regulation of trans-synaptic signalling) as well as WNT signalling. The congruence of cluster 2 effects across both cohorts suggests that these specific populations of astrocytes consistently associate with a prior history of high adversity exposure across psychiatric disorders in the human PFC.

Finally, we performed differential expression analysis in the astrocytes robustly assigned to each cortical layer. Relative to the low-adversity-cases, the high-adversity-cases had increased ERMN and ERBB3 in the white matter, APBB3 in layer 2, and decreased PCDH9 in layer 5 (Supplementary Figure 7, Supplementary Data 10). Neither group had any significant DEGs compared to the low-adversity-control group. Together, this demonstrates that specific cortical astrocyte population have phenotype that associates with prior exposure to high levels of adversity in the context of psychiatric disorders.

### Females with psychiatric diagnosis and high adversity exposure demonstrate more pronounced astrocyte-specific gene expression changes

Transcriptomic profiling across multiple psychiatric disorders demonstrates notable transcriptomic differences in the PFC of males and females^37,38^, which are relevant at the single-cell resolution in the PFC of these disorders^14^. To explore whether this pertains to astrocytes, we performed a sub-analysis of differential gene expression in males and females separately (55 males, 32 females; Figure 4a). Between the adversity-stratified cases, autosomal DEGs were almost exclusively observed in the female clusters (98%), and separating the analysis by sex increased the number of DEGs detected in both males and females. These female-specific DEGs reflected the DEGs before stratification by sex, with striking differences in metabolic cluster 2 (increasing from 40 to 94 DEGs) and homeostatic cluster 3 (increasing from 0 to 351 DEGs; Figure 4a, Supplementary Data 11). To further support sex-biased effects, we performed rank-rank hypergeometric overlap between male and female DEGs low vs high-adversity exposed cases. There were very weak positive correlations between the ranks of male DEGs and female DEGs at both low-resolution (protoplasmic/fibrous) and high-resolution (cluster) levels (ρ < 0.09; Figure 4b; Supplementary Figure 8). The one exception was cluster 0 which was strongly correlated (ρ = 0.49). Reflective of prior bulk transcriptome studies^37^, diagnosis-low-stress and diagnosis-high-stress groups both demonstrated moderate to strong negative correlation between male DEGs and female DEGs at high resolutions, compared to control-low-stress (ρ = 0.36-0.41; Figure 4b). For both males and females, the metabolic cluster 2 (*SLC1A2*) and homeostatic cluster 3 (*LSAMP*) contributed almost all DEGs in these comparisons (Supplementary Figure 9). These results indicate that female cases of psychiatric disorders may be driving astrocyte differential gene expression associated with high adversity exposure.

**Figure 4:**
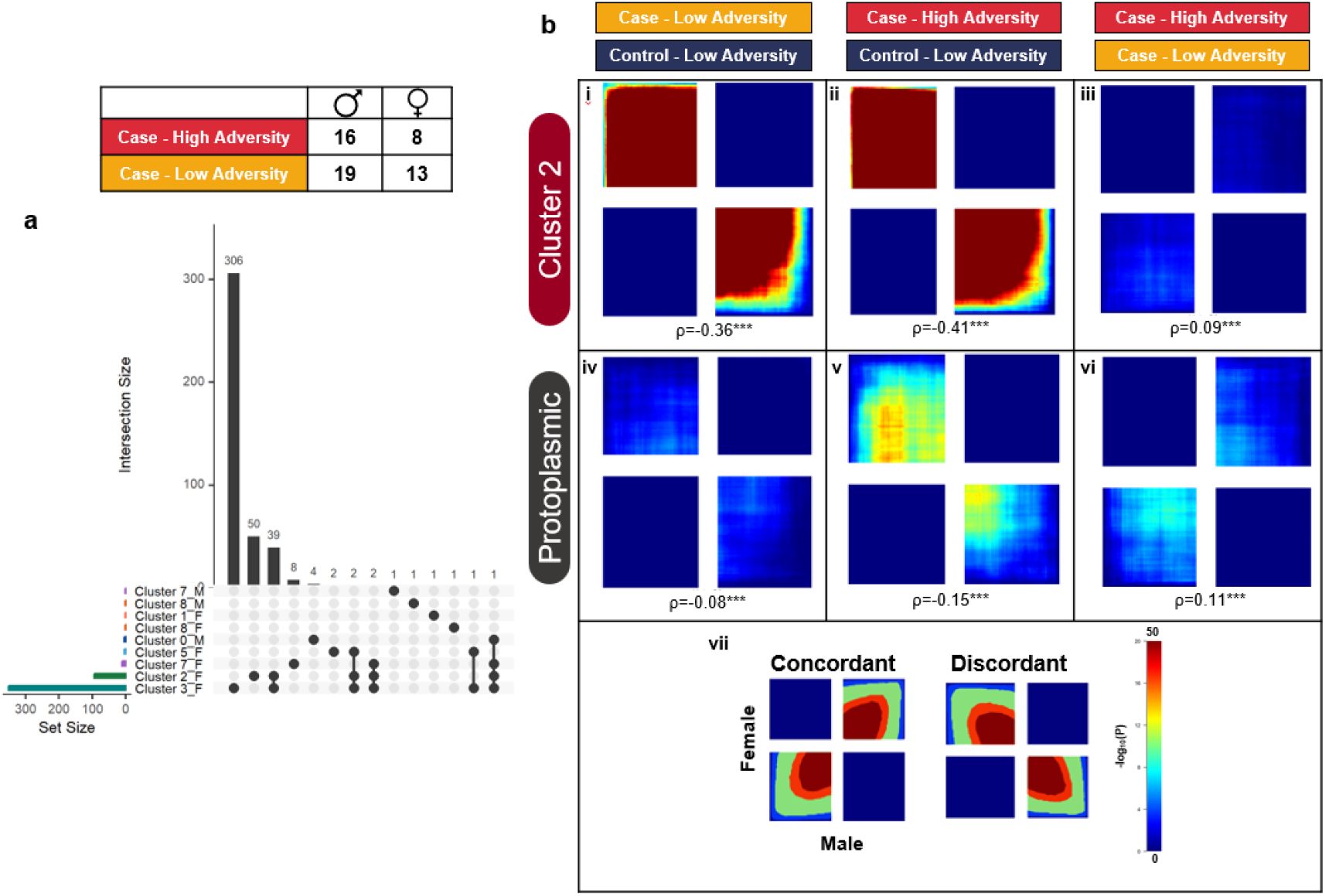
Sex demonstrates discordant case-control effects in homeostatic astrocytes, females demonstrate stronger transcriptomic associations with adversity. **(a)** UpSet plot depicting the differentially expressed genes between high- and low-adversity cases (P_adj_<0.1). Set size indicates number of genes meeting threshold for each cluster and dots connected by a line indicate common genes. F indicates data derived from females. **(b)** Rank-rank hypergeometric overlap between differentially expressed genes stratified by sex. Panel i-iii denote cluster 2 comparisons, iv-vi denote broad protoplasmic comparisons, panel vii denotes shared dimensions, including scale range, explanation of quadrants and axes. Heatmap indicates direction of shared genes, concordant indicates that DEGs are consistently ranked between males and females out of all genes and discordant the reciprocal trend. i and iv denote no stress cases-control, ii and v denote stress cases-control, and iii, iv denote stress cases-no stress cases.

### Timing of adversity exposure associates with transcriptomic and morphological changes to astrocytes

Given that early life adversity disproportionately contributes to psychiatric disorder risk^39^, we explored whether the timing of adversity exposure influenced gene expression. We performed a second sub-analysis on the cohort based on the timing of first occurrence of severe adversity exposure in the medical records (childhood (<12, n=6), adolescence (12-18, n=6), early adulthood (18-25, n=5), adulthood (>25, n=7) and low adversity. We performed differential expression analysis treating adversity timing as a continuous outcome in the limma analysis (one indicates no adversity and five indicates earliest timepoint of adversity). In total, five genes in cluster 2 and two in cluster 3 were significantly associated with adversity timing (P_adj_<0.1, Supplementary Data 12). These related to synapse adhesion (e.g. *LSAMP, CADM2*; Figure 5a/b) and interestingly, *PCDH9*, which was decreased in layer V astrocytes associated with high adversity, also associated with earlier adversity timings. The DEG profile of adversity timings for cluster 2 were similarly moderately associated with those identified in the replication cohort (ρ=0.25, Supplementary Figure 10a).

**Figure 5:**
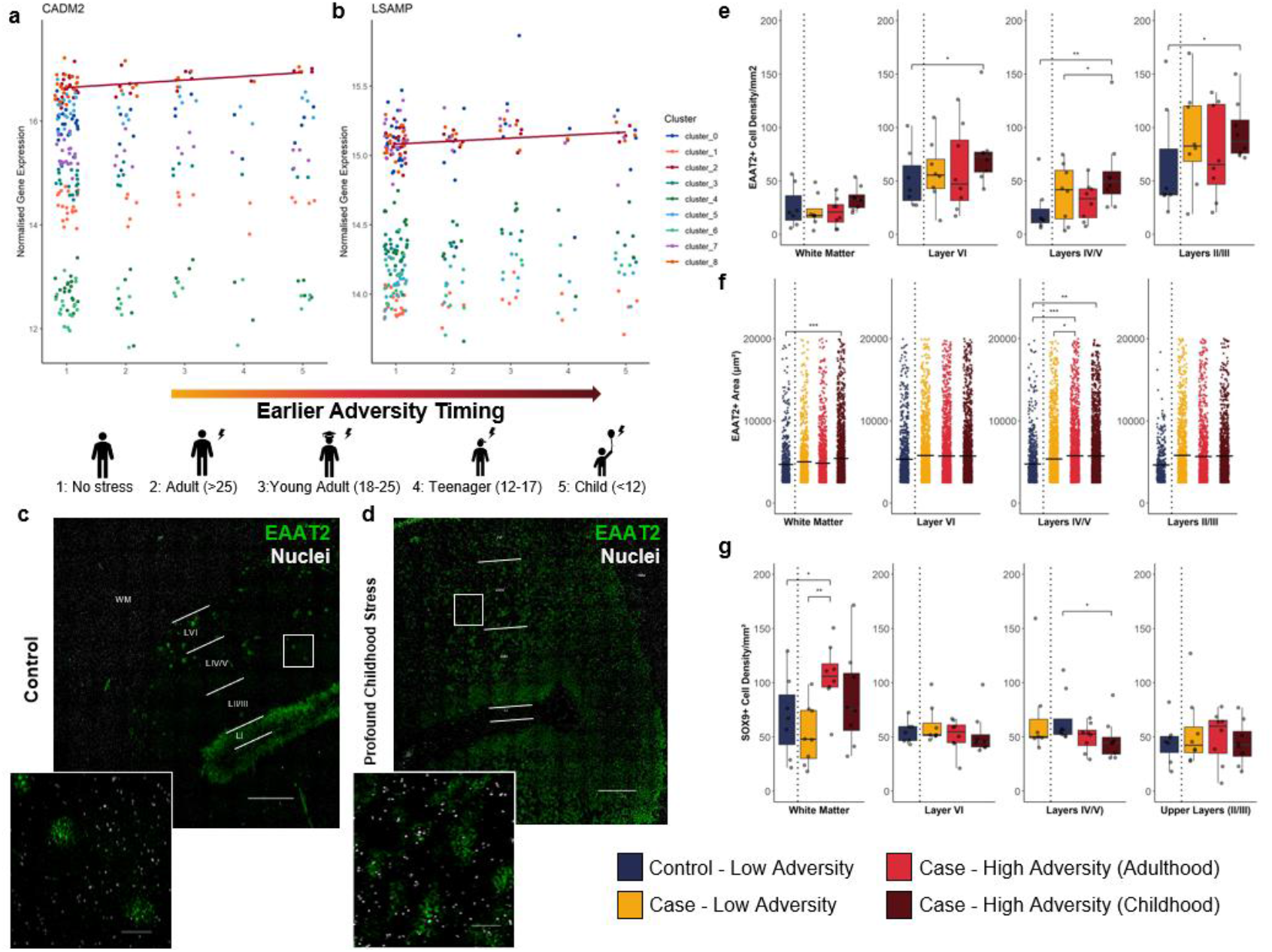
Earlier exposure to high adversity load associates with greater alterations to astrocytes across transcriptomic and morphological measures. **(a-b)** Scatterplots for genes associated with adversity timing, based on key timing of exposure (5: childhood (<12), 4: adolescence [12-18], 3: early adulthood [18-25], 2: adulthood [25+] and 1: low adversity before diagnosis). Normalised expression was corrected for visualisation using batch and covariates via limma removeBatchEffects (left to right: CADM2, LSAMP). Point/line colour indicates cluster. Line indicates significant association of given cluster with adversity timing (P_adj_<0.05). **(d)** Representative image of EAAT2 immunohistochemistry with representative automated segmentation used for morphology analysis. Scale bar = 1mm. **(e)** Density of EAAT2+ cells per mm^2^ by adversity stratification and cortical layer. Box plots indicate median and quartiles. Significance calculated using linear regression on the log value correcting for covariates. *P<0.05,**P<0.01. **(f)** Area of EAAT2+ cells. Individual cell values are plotted. Line indicates mean. Significance calculated using linear regression on inverse value corrected for covariates as well as derived individual. P conservatively adjusted for number of cells using Bonferroni correction (n tests = number of cells). *P<0.05,**P<0.01. *P_adj_<0.05, **P_adj_<0.01, ***P_adj_<0.001. **(g)** Density of SOX9+ cells per mm^2^ by adversity stratification and cortical layer. Box plots indicate median and quartiles. Significance calculated using linear regression on the log value correcting for covariates.

To support this sub-analysis, we quantified the morphology and density of astrocytes expressing the glutamate transporter EAAT2 (encoded by *SLC1A2*) using immunohistochemistry on whole coronal sections from a well-matched and previously validated subset of this cohort (Figure 5c/d, cohort outlined in Supplementary Figure 3)^40^. We chose EAAT2 given that cluster 2, which exhibited the adversity-associated effects across all analyses, was enriched for the transcripts of encoding this protein and hence best captured by quantification of this marker. This cluster also had relatively low expression of other common markers such as GFAP^41,42^, further incentivising this astrocyte-enriched protein. We quantified the morphology of the domain of 21,794 EAAT2+ astrocytes based on laminar localisation (layer II/III, layer IV/V, layer VI, and white matter). The density of EAAT2+ astrocytes nominally increased in high-childhood-adversity-cases (<12 years) compared to low-adversity-controls across all layers (11-39%; VI-II; P=0.002-0.016), and low-adversity-cases, specifically in layers (IV-V) of the grey matter (20%; P=0.035), with only the childhood-control comparison in layers IV-V surviving testing for multiple comparisons (P_adj_=0.014, Figure 5e). There was also an increase in the territory covered by EAAT2+ astrocytes across all high stress cases in layers IV-V (adulthood: +21%, P_adj_=2.99×10^−5^, childhood: +21%, P_adj_=2.43×10^−4^) as well as an increase in the white matter (+15%; P_adj_=2.31×10^−11^) of those with childhood adversity (Figure 5f). Complementary morphology measurements (perimeter, circularity, eccentricity, and max width) bared similar differences (Supplementary Figure 10b). In further support that EAAT2+ astrocytes were distinctly implicated, we also quantified density with the total astrocyte marker SOX9^43^ and found minimal changes in density between groups in the grey matter, with only decreases in the layers IV-V in cases with childhood adversity, compared to those without (Figure 5g, Supplementary Figure 10c). This finding corroborates our transcriptomic evidence that there are detectable and persistent differences to astrocytes in human brains that have previously experienced high adversity. Importantly, these effects are not observed ubiquitously across cortical astrocyte populations nor spatial domains of the cortex, with astrocytes identified by high expression of *SLC1A2* (EAAT2) in the grey matter disproportionately impacted.

### Human astrocyte neurotransmission is sensitive to chronic glucocorticoid activation

Finally, it is unclear to what extent human astrocyte function is impacted by a direct biological stress response and by indirect effects from the surrounding environment. To directly test the responsivity of astrocytes to stress, we differentiated human astrocytes (iAsts) using lentiviral delivery of the transcription factors *SOX9* and *NFIB* to neural progenitors derived from the female hESC line (H9) (Figure 6b i-iv)^44,45^. Synthesised iAsts expressed key astrocyte marker RNA (qPCR; Figure 6c) and proteins (immunocytochemistry; Figure 6b v), could effectively clear glutamate from media, and had robust calcium responses to agonists including glutamate and ATP (Figure 7; Supplementary Figure 12). They were also largely pure; 95% expressed *GLUL*, 94% *SLC1A3*, and 93% *SOX9* (Figure 7a-c, Supplementary Figure 13c-e). Importantly, the expression of glutamate receptor (NMDA and AMPA receptors) subunits largely resembled proportions observed in the snRNAseq dataset, except for the AMPA receptor subunit *GRIA3*, NMDA receptor subunit *GRIN2B*, and metabotropic glutamate receptor *GRM3* (Figure 6c). To mimic acute exposure hormonal stress, iAsts were exposed to the glucocorticoid receptor agonist dexamethasone (100nM)^46^. The stress paradigm was conducted for durations of acute (12 hrs) to chronic (seven days), at which point the culture could not be effectively maintained due to increased proliferation. This stress paradigm caused a dramatic increase in the expression of glucocorticoid-induced transcripts *TSC22D3* and *FKBP5* and cells immunopositive for FKBP51 (the protein encoded by *FKBP5*) by three days of treatment, with almost all cells immunopositive by seven days of treatment (Figure 6d).

**Figure 6:**
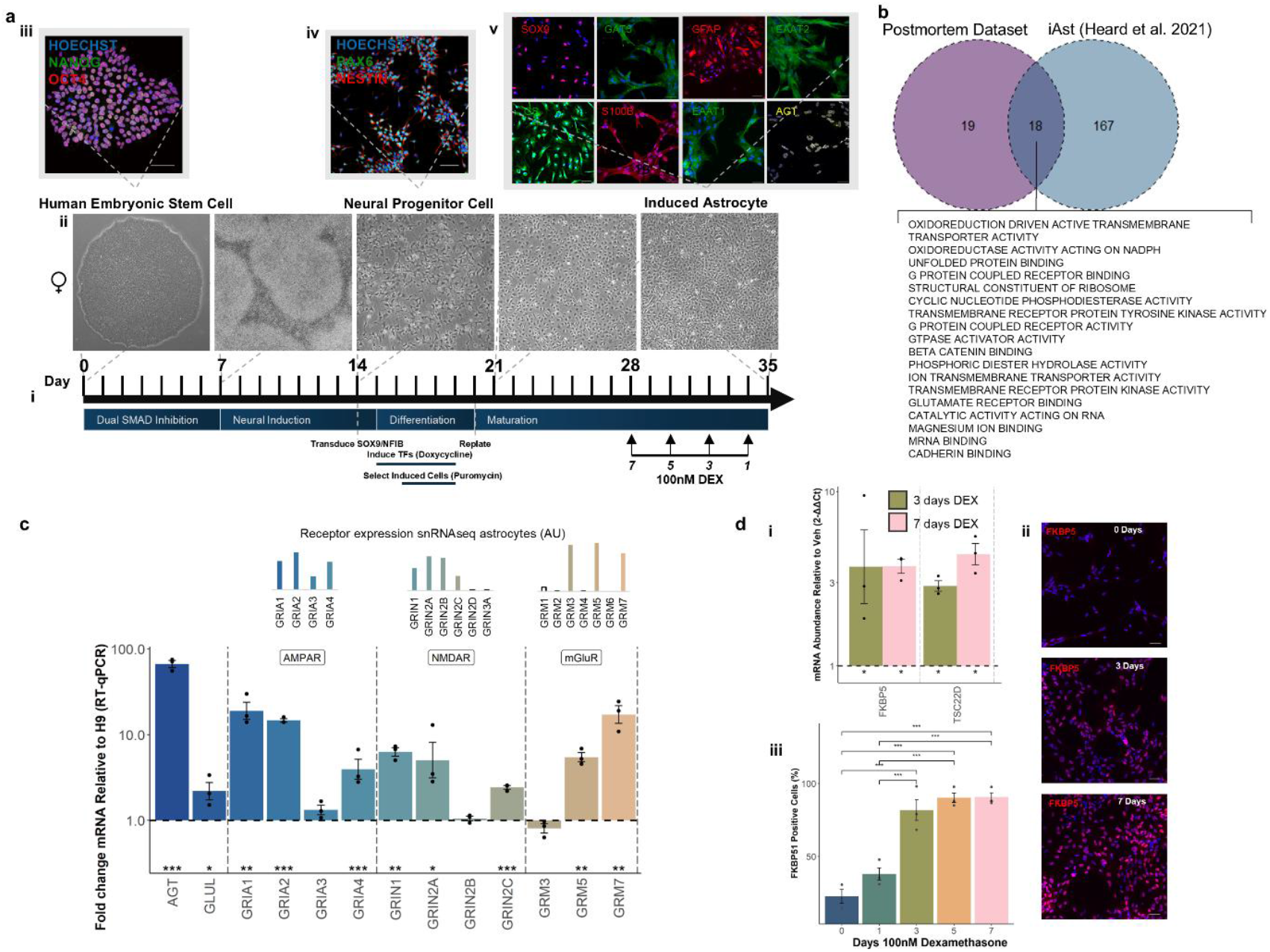
Human pluripotent stem cell-derived astrocytes can model specific dimensions of human astrocyte sensitivity to glucocorticoids. **(a)** Differentiation protocol used to derive astrocytes and expose them to dexamethasone: i) Timeline of differentiation; ii) Representative brightfield images of differentiation protocol; iii-v) Representative immunocytochemistry images from stages of differentiation validating expression of cell type markers. Scale bar = 50µm. **(b)** Venn diagram of significant terms returned from all clusters and iPSCS using the Molecular Function GO database. **(c)** Fold change expression of genes in H9-derived iAsts compared to H9 stem cells. Astrocyte maturity markers (*AGT* and *GLUL*) and subunits of ionotropic (NMDA and AMPA) and metabotropic (*GRM3/5/7*) glutamate receptors. Bars represent mean value ± 1SD. Graphs above represent relative expression of subunits in postmortem dataset in arbitrary units (AU) **(d)** Validation of response to glucocorticoid paradigm: i) Fold change expression of glucocorticoid-responsive genes at 3 and 7 days of exposure relative to vehicle; ii) Representative immunocytocheemistry images with normalised FKBP51 staining at 0, 3, and 7 days of exposure; iii) Percentage of FKBP51 positive cells across timepoints. Calculated using ANOVA with post-hoc TukeyHSD. Bars represent mean value ± 1SD. *P<0.05, **P<0.01, ***P<0.001. For all experiments, n=(3 biological replicates [different differentiations from hPSC] x 2 technical replicates [different well assayed]).

To identify specific functions to profile with this model, we used GSEA on an existing dataset of cortisol-treated astrocytes derived from female donors^47^. This identified 18 altered molecular functions shared with functions associated with adversity in the postmortem cohort (Figure 3e, Figure 6b). Of these functions, 12 were common to cluster 2, including G Protein-Coupled Receptor Binding and Glutamate Receptor Binding, both modulators of glutamate signalling (Figure 6b). As a result, we chose to focus profiling the impact of the stress paradigm on glutamate-associated functions of astrocytes.

First, we considered the role of astrocytes in clearing glutamate. While there was no significant effect of dexamethasone treatment on the number of EAAT1, EAAT2, or glutamine synthetase-positive astrocytes (Figure 7a-c), we did see a transient decrease in glutamate clearance after three days of ‘stress’, which was returned to untreated levels by seven days (Figure 7d). We also employed whole-well ratiometric Ca2+ imaging, to measure the maximal responses of treated iAsts to glutamate, NMDA and AMPA agonism. After seven days of exposure, we observed that the maximum peak response of the iAsts to both glutamate and NMDA increased, relative to vehicle (8.3% increase, P=0.004; 8.1% increase, P=0.0047, respectively, Figure 7e-f). There were no significant differences in the Ca2+ response to AMPA following treatment with dexamethasone (Figure 7g).

We then quantified RNA expression of receptor subunits involved in coordinating direct Ca2+ responses, including NMDA, AMPA, and metabotropic receptors (mGluR3/5/7) (Figure 7h). We identified alterations to both NMDA and AMPA receptor subunits, with changes across all AMPA subunits (GRIA1/GRIA2 up, GRIA3/GRIA4 down), and NMDA subunits GRIN1 (up) and GRIN2A (down). There were also modest decreases in the metabotropic receptors GRM3/GRM5, although the former was expressed at a low level. Notably, several of these transcript level effects were conserved with the study used to perform GSEA, including decreased GRIA2/GRIA3/GRIA4 and GRIN2A^52^.

To validate these findings, we used data from a separate model of human astrocytes. Astrocytes were isolated from cerebral organoids derived from iPSCs from three separate individuals, exposed to either 100 nM dexamethasone or vehicle for three days, and RNA was sequenced (oAsts). Consistent with our initial iAst model, in the oAsts treated with dexamethasone, we found increased SLC1A3 (log2FC=1.39, Padj=0.009) and GRIA1 (log2FC=3.22, Padj<0.0001), alongside decreased GRIA3 (log2FC=-0.94, Padj=0.01), and GRIN2A (log2FC=-2.50, Padj<0.0001). In further support, GSEA using the Biological Processes database showed that the most significantly enriched function was decreased regulation of trans-synaptic signalling (NES=-1.62, Padj=0.01). These findings consistently indicate that astrocyte functions in neurotransmission are actively shaped by activation of the glucocorticoid-mediated stress system, and consistent with astrocytic alterations we observed in individuals with psychiatric disorders with a history of high adversity exposure.

## Discussion

This study profiled the transcriptome of >100,000 astrocytes from 87 individuals and morphology of >20,000 astrocytes and considered these with spatial resolution. We identified that in individuals with a major psychiatric disorder, a prior history of high adversity exposure was associated with identifiable changes to the transcriptomic landscape of astrocytes. These changes were pinpointed to a population of grey matter astrocytes demarcated by a strong neurotransmitter reuptake profile. We validated that these functions are directly shaped by hormonal systems central to the stress response in hPSC-derived astrocytes from two- and three-dimensional cultures exposed to dexamethasone, a potent glucocorticoid receptor agonist. Together, our data consistently indicated that high adversity exposure associates with lasting changes to the grey matter astrocytes, particularly those performing glutamate-related functions.

We first integrated multiple high-resolution transcriptomic techniques to better understand the diversity of astrocytes in the human OFC. This basis of diversity was then used to explore the lasting impacts of adversity exposure in the context of psychiatric disorders. This implicated a population of astrocytes demarcated by particularly high expression of glutamate transporters (*SLC1A2/3*) in delineating individuals with high adversity exposure from those with lower adversity exposure. Prior snRNAseq approaches investigating psychiatric disorders have identified limited associations between astrocytes and either schizophrenia or major depression^10,13,48,49^. A caveat of these approaches is that astrocytes are regularly conglomerated into a single population, while neuronal cell types are delineated into several populations, based on laminar and molecular phenotypes. While examining broad cell types is crucial to narrowing the focus of the immense amount of data presented by high-resolution sequencing techniques, it likely masks heterogeneity existing in non-neuronal cell populations^50-52^. Our high-resolution approach, enabled by the analysis of >110,000 astrocyte transcriptomes (∼2.5-14 times more than in prior human studies in the cortex^26^, and comparable to a recent large-scale hippocampus atlas^27^) represents one of the most comprehensive characterisations of human astrocytes to date. We also cross-validated this with a second cohort of PTSD to support that these effects are seen across a range of psychiatric disorders. This resolution revealed that both the transcriptomic and spatial diversity of astrocytes are relevant to psychopathology.

There have been indictors that astrocytes are heterogeneously associated with human psychopathology. One study that performed qPCR on microdissected layers of the human anterior cingulate gyrus identified that several astrocyte markers, including the glutamate transporter EAAT2 and metabolising enzyme glutamine synthetase (GS), were selectively reduced in the deeper layers of the cortex (IV-VI), but not layers I-III nor in the white matter of individuals with schizophrenia^53^. Indeed, when we considered the astrocyte population at a bulk level (protoplasmic/fibrous), we observed no DEGs between the adversity-stratified cases and only 10 DEGs between cases and controls. Our high-resolution analysis of astrocytes strengthens evidence that astrocyte diversity is relevant to adversity-associated pathologies.

In non-human species, there is evidence of the transcriptional impact of adversity on astrocytes, even at a broad cell-type level. A recent snRNAseq and spatial analysis of the DLPFC from female macaque primates identified that astrocytes contributed the most DEGs associated with social stress^54^. In line with the present study, they identified that cell adhesion and glutamatergic synapse activity were over-represented terms associated with these DEGs, including decreased glutamine synthetase (*GLUL*), increased metabotropic (*GRM5*) and ionotropic (*GRIA1, GRIK2*) glutamate receptors in females. Female rodents have demonstrated divergent responses to cortical astrocytes, with only females demonstrating increased cell area in response to chronic restraint stress^55^. Estrogen can also increase EAAT2 expression and glutamate uptake in primary astrocyte culture^56^. Sex-specific temporal cues are important drivers of astrocyte maturation, with astrocytes in male rodents reaching maturity faster than in females^57^, suggesting that the risk window for astrocyte sensitivity may diverge based on sex, and that sex hormones may influence this. Indeed, the hypothesis that adversity occurring early in life when the brain is vulnerable is crucial to disorder onset has existed for decades,^58-60^ and sex is likely a strong mediator of these changes^61^. Another study profiling the transcriptome of cortical astrocytes derived from male rodents exposed to chronic variable stress identified upregulation of *GRIA2* and *GRIA1*, in agreement with our glucocorticoid-treated iAsts^62^. The potential of these models to capture these changes at the bulk transcriptome level may also be partly due to their capacity to control the genetic and environmental conditions within the sample, something not possible in human studies.

The gene ontology terms emerging from this study indicate that the coordination (signalling and adhesion) of the synapse is a key system delineating high-adversity-cases of psychiatric disorders. Reduced cortical synapses on excitatory neurons are one of the most well-documented phenomena associated with early life adversity in rodents^11,63^. We have previously demonstrated that these effects are likely translationally robust; in the same cohort used for morphological analysis we previously identified reduced dendritic spines in cases with profound childhood adversity^40^, and that mature spine density is negatively correlated with stress modulators such as *FKBP5*^64^. A recent study demonstrated that cortisol activation of the glucocorticoid receptor directly mediated the pruning of excitatory synapses via astrocytes in both rodents and human-derived cortical organoids^65^. Importantly, this effect also attenuated behavioural symptoms associated with early social deprivation. Another study co-culturing mouse glia with human-derived neurons demonstrated that astrocytes coordinate the expression of several neuronal synaptic genes associated with schizophrenia^66^. They also noted that physical contact with astrocytes drove the expression of these genes, including *NRXN1*, which was increased in association with stress in cluster 2. Recent snRNAseq evidence also suggests that astrocyte synapse functions are altered in schizophrenia and further moderated by ageing^13^. This evidence supports that the modulation of synapses by astrocytes is a central system relevant to psychiatric disorders and that these systems are directly shaped by glucocorticoid signalling. We demonstrated across two separate human-derived astrocyte models (iAsts and oAsts) that several of these functions are directly shaped by glucocorticoid activation, and that AMPA receptor subunits *GRIA1* and *GRIA3* are impacted consistently. Targeting astrocytes may, therefore be a promising target for novel therapeutics and may be able to leverage these systems.

Finally, some key limitations of this study should be acknowledged. We used snRNAseq, not single cell RNA sequencing (scRNAseq), which expectedly limits the detection of transcripts related to ribosomes and mitochondria but also has relevance to glial cell states such as disease-associated microglia^67^. For most genes, however, results are largely concordant with single-cell approaches^68,69^. Another technical limitation was the inference of the snRNAseq onto the spatial transcriptomic domains. While we found parallel trends between imputed laminar distributions and immunohistology, targeted sequencing methods may more faithfully capture transcriptomic changes associated with laminar populations (e.g. targeted laser capture paired with scRNAseq). Another limitation was the sample size, which limited power in exploratory analyses (e.g. sex-specific and adversity-timing analyses). Despite this, this postmortem study is currently the largest to date. Another future consideration would be the inclusion of controls with high adversity exposure if donor tissue becomes available. However, out of the 33 healthy controls, only one had a notable history of adversity, of particular note given that high levels of adversity are such a strong risk factor for later developing a psychiatric disorder.

Of note, iAsts and oAsts most likely resemble late foetal astrocytes and thus have limited transcriptomic similarity to mature human astrocytes (mean donor age in the present study > 50 years). The cellular environment around astrocytes is also likely not accurately represented in *in vivo* models^70-72^. This may limit the expression of astrocyte adhesion molecules^66^, a key hit arising from the postmortem tissue work. Another consideration is that assays were performed in relatively acute conditions (up to seven days), without a recovery window. Consequently, this model likely does not reflect changes lasting beyond the removal of the stressor, which would provide a useful direction for follow up. We also focused on the glutamatergic functions of astrocytes given that they are conserved hits between the datasets. However, functions associated with RNA and ribosomes were also conserved and may be promising targets for further exploration. Alterations to both RNA and ribosome processing have been strongly identified in studies of the human cortex in schizophrenia^73,74^ and glucocorticoids directly regulate these processes^75^. Given that iAsts proliferated upon exposure to DEX, we opted not to focus on these biosynthesis functions, particularly as evidence of astrocyte proliferation associated with brain disorders is limited^76-78^. Finally, astrocyte calcium signals were measured by bulk signal (per well). This was to allow for multiple endpoints to be assayed in near tandem and in replication (not feasible in typical setups). It is worth noting that this signal may be in part associated with underlying changes to other systems, such as gap function coupling^79^ and may be more reflective of astrocyte communication deficits. However, probing with multiple targeted agonists to different effect supports at least some of this effect is due to agonist activity.

In summary, this study demonstrates that astrocytes are a highly diverse cell population and delineable across molecular, spatial, and morphological measurements when stratifying psychiatric disorder cases based on adversity history. We identified that astrocytic coordination of the synaptic environment in the grey matter is a convergent system capable of delineating ‘stressed’ astrocytes from ‘unstressed’ astrocytes in humans, including in multi-transcriptomic, histological, and in vitro modalities. Our findings in astrocytes support that studies of psychiatric disorders at high resolution should aim to better reflect the clinical heterogeneity core to these disorders, for example adversity exposure, as this will likely be essential to improving treatment specificity, and thus efficacy.

## Methods

### Postmortem human brain samples

Ethical approval for the study of postmortem human samples was obtained from the Human Ethics Committee at the University of Wollongong (HE 2020/308). The history of adverse events (childhood (<12 years), adolescence [12-18 years], early adulthood [18-25 years], adulthood [25+ years] and low adversity before diagnosis) and psychiatric diagnoses (schizophrenia, schizoaffective disorder, depression, bipolar disorder) were extracted from extensive medical records of 92 adults (58 males, 34 females), as previously described^40,64^. Briefly, individuals were stratified based on whether they had a diagnosis of a severe psychiatric disorder, and whether they had been exposed to high levels of adversity before disorder onset (32 no diagnosis, 32 with a diagnosis and with low adversity exposure, 28 with a diagnosis and with high adversity exposure). High adversity was considered an event, series of events, or set of circumstances that was physically or emotionally harmful or threatening and had lasting adverse effects on the individual’s functioning and physical, social, or emotional well-being (e.g. physical/emotional abuse or neglect). From this cohort, fresh frozen tissue from the 3^rd^ 8-10mm coronal section of Brodmann Area 11 (OFC) was used for all experiments. For single nucleus RNA sequencing, 87 samples produced high-quality libraries (one individual had low data quality and four had over 50% of nuclei were clustered in a single cluster) consisting of 32 low-adversity-controls, 31 low-adversity-cases, 24 high-adversity-cases^20^. One sample was omitted from analysis related to psychopathology due to having a profound history of adversity, but no noted psychiatric diagnosis. A subset of these individuals with histologically sound tissue structure were also processed for spatial transcriptomics (n=13). Finally, an overlapping subset (n=32, 27 individuals common between approaches), in which we have previously identified neuronal associations with childhood adversity exposure^40^, was used to explore the morphology of astrocytes. Groups had non-significant distributions of sex (complete cohort: χ^2^=0.80, P=0.67, cohort used for snRNAseq: χ^2^=0.08, P=0.96) diagnoses (complete: χ^2^=1.83, P=0.61, snRNAseq: χ^2^=2.75, P=0.43), according to a chi-squared test. A schematic breakdown of the cohort, as well as key cohort demographics (diagnosis, age, sex, postmortem interval, RNA integrity, and brain pH) are detailed in Supplementary Figure 1.

### Single nucleus RNA sequencing

#### Sequencing, preprocessing, and broad cell type annotation

Nuclei were extracted and processed from fresh frozen postmortem brain tissue as described previously^20^, with 10,000 nuclei per sample as target recovery. Briefly, samples were sequenced using the NovaSeq 6000 System and reads were aligned using Cell Ranger (Version 6.0.1) ‘count’ pipeline (genome build GRCh38, Ensembl 98). Reads per cell were downsampled to the 75% quantile and integrated in Python using Scanpy (Version 1.7.1)^80^. Preprocessing steps included filtering genes in <500 nuclei and cells with <500 counts, <300 genes, and/or >15% mitochondrial genes, doublet removal^42^, and data normalisation and log-transformation using sctransform (Version 0.3.2)^81^. The Leiden clustering algorithm^82^ based on highly variable genes was used to cluster nuclei (resolution 1.0). Cell types were assigned using a label transfer algorithm to infer primary cell types in clusters and high-resolution clusters were further manually annotated using marker genes as previously described^20^. Over 800,000 sequenced nuclei underwent quality control, clustering, and cell-type labelling^20,64^. a cluster of 113,140 nuclei labelled as astrocytes were isolated from this larger dataset for further analysis, using Scanpy for data processing (Version 1.9.6).

We then segregated and performed targeted re-clustering on these cells to explore heterogeneity at a higher resolution, identifying an initial 12 astrocyte clusters (Supplementary Figure 2). We removed two clusters given that one contained low-quality reads (high mitochondrial counts), and another was derived predominantly from two individuals.

#### Clustering astrocytes

In the astrocyte subset, highly variable genes and connectivity of neighbours were re-calculated using the same process as above. Dimensional reduction via uniform manifold approximation and projection (UMAP) was subsequently performed for visualisation. Appropriate clustering of astrocytes was determined using the Leiden algorithm. Stability analysis was conducted at resolutions 0.2-1.2 using two bootstrapping approaches. In the first, 90% of cells were subsampled and Leiden clustering with random seeding was applied across resolutions over 20 iterations. Stability per cluster was calculated as the mean value of all cells in that cluster being sorted into the same cluster. While the most stable clustering was at 0.2, 0.8 was selected due to its high stability, relative to other resolutions, capacity to replicate clusters from prior studies (i.e. cluster 1), and ability to capture low cell count clusters (i.e. cluster 9; n=158 cells [0.14% of astrocytes]). To further support this, we used a second more stringent bootstrapping approach. Twenty iterations of Leiden clustering were applied to the whole dataset to determine baseline stability. A second round of 20 iterations was performed, removing the cells equivalent to one cluster at that resolution, The difference in stability between these two iterative rounds was then determined. This further identified 0.8 as the most stable resolution for clusteriong. Marker genes per predicted cluster were determined using the rank_genes_groups function in Scanpy (method=‘Wilcoxon’), in accordance with recommendations with a recent benchmarking paper for marker gene selection^83^.

#### Differential expression analysis

Differential expression was conducted using limma (version 3.18, R)^84^. First, pseudobulk counts (the sum of raw counts per cluster, per individual) were calculated using ADPBulk (Python), and variance partitioning was performed on pseudobulk counts from all individuals using VariancePartition (R, version 1.28.9) fitting for measured covariates complete across the cohort (Batch, Sex, Age, Postmortem Interval, RNA Integrity, Brain pH, Adversity Stratification)^85^. Variance partitioning identified strong effects of processing batch effects, high association of sex-specific genes (*UTY, XIST*) with sex, brain pH with ribosomal genes (*RPS27, RPL35*) and a significant proportion of variance unexplained by captured cohort demographics (Supplementary Figure 6). To correct for unexplained variance, the pseudobulk counts were first transformed using the limma voom transformation. The removeBatchEffect function (limma) was then used to account for all covariates mentioned above. Principal component analysis was then performed on the corrected counts and the resulting first principal component was included as a covariate in differential expression analysis to account for the high degree of unexplained variance. The final model used for differential expression was (∼ library batch + sex + age + postmortem interval + RNA integrity + brain pH and PC1 (unexplained variance)). All groups were included in the model, and pairwise comparisons between each of the groups was used to calculate DEGs. For sex-stratified analysis, the same approach was used, omitting the ‘sex’ variable from the model (∼ library batch + age + postmortem interval + RNA integrity + brain pH and PC1). Extremely small pseudobulk libraries <2000 reads were omitted. Analysis was conducted on robustly expressed genes with counts per million (CPM) >10 in at least 25% of the cohort, reflective of ∼80% of the smallest group. DEGs were calculated between the cases (cases-high-adversity vs cases-low-adversity) as well as healthy controls (cases-high-adversity vs cases-low-adversity). Significance was considered any gene surviving a false discovery rate of 10% (P_adj_<0.1, Benjamini-Hochberg correction). We omitted cluster 9 from differential expression analysis due to low number of cells.

#### Gene set enrichment analysis

Gene set enrichment analysis was performed using either cluster markers or DEGs using fGSEA (version 3.18, R). For cluster markers, rank was calculated as the gene score from the Wilcoxon test. For DEGs, rank was calculated as in a previous report^14^ (-log10(nominal P) * sign(log2FoldChange)) and gene ontology databases (Biological Processes, Molecular Function, Cellular Compartment) were accessed via msigdb (msigdbr version 7.5.1, R). For visualisation, terms were pruned based on semantic similarity, using GO.db (version 3.16.0, R) to access the GO directed graphs. Terms were only kept for visualisation if they had no nominally significant child terms (P<0.05). Full, unpruned lists are presented in Supplementary Data 8. Significance was set at P_adj_<0.05.

#### Gene-disease association (GeDiPNet 2023)

The online tool EnrichR was used to access the GeDiPNet 2023 database. Unranked significant DEG lists (P_adj_<0.1) were input and resulting gene ontology terms were sorted according to combined score. Significant terms were considered those with P_adj_<0.05.

#### Rank-rank hypergeometric overlap

Threshold-free comparison between the DEGs of various analyses was performed using rank-rank hypergeometric overlap (RRHO) analysis available via the RRHO2^86^ package. Genes were scored using scored as for GSEA (-log10(nominal P) * sign(log2FoldChange)). Scored gene lists were initialised using the RRHO_initialise function (method = ‘hyper’, log10.ind=TRUE) and visualised using a heatmap. The direction and strength of DEG correlation was further calculated using Spearman’s correlation between each ranked gene list.

### Validation Single Cell Dataset

Published snRNA-seq data from DLPC snRNA-seq cohort by Chatzinakos et al. was used to replicate adversity findings. It was composed of 11 individuals diagnosed with PTSD and 11 controls (362,996 nuclei, 34,504 astrocytes). A detailed pipeline of all processing steps is provided by Chatzinakos et al.^10^. Both post-processed, normalised and integrated count matrices were exported and were imported in Seurat for further processing.

#### Astrocyte Reclustering

Astrocytes from the validation cohort were reclustered based on the identified astrocyte clusters of the snRNAseq dataset. Variable features were detected with Seurat::FindVariableFeatures, scaled the data with Seurat::ScaleData and performed PCA with Seurat::RunPCA to detect 100 PCs, which was used as embeddings to create the tSNE reduction with Seurat::RunTSNE. The Seurat::FindTransferAnchors was used with the primary dataset as reference the replication dataset as the query to project the clustering and applied Seurat::TransferData to extract information on the number of nuclei from the replication snRNAseq dataset that are allocated in the cell types/subtypes of the primary snRNAseq dataset.

#### Differential gene expression within the various astrocyte clusters

To calculate pseudobulk (sum of counts) expression matrices for cell-clusters and /or annotated cell-types cluster, the raw unique molecule identifiers (UMI) counts of the corresponding single-cells were summed to estimate one value per cell per individual for each gene. The summed counts were converted to counts per million (cpm) function in edgeR 2. Genes with cpm > 2 for at least 66% of the total sample size were selected based on the edgeR::filterByExpr(). In addition, TMM was used to estimate normalization factors that account for library size variation between samples. The normalization factors were then considered by limma::voom that was used to estimate the mean-variance relationship of the log-counts and to generate a precision weight for each observation. Finally, the voom-normalized feature counts (on the log2 scale) were used for linear modelling. Differential gene expression was based on trauma load using the limma package^84^. Trauma was quantified based on exposure to traumatic events; 0 indicates no trauma exposure (controls), 1 adulthood trauma, and 2 childhood trauma. Given that trauma exposure is core to PTSD diagnosis, a diagnosis group without adversity was not feasible. As a result, a linear regression fit for each gene, using trauma timing from the above three groupings as the outcome. Further details on the linear regression models can be found in prior publication^10^.

### Spatial transcriptomics

#### Library preparation (Visium)

Fresh frozen orbitofrontal cortex (BA11) sections (10µm thick) spanning all cortical layers were mounted onto capture areas on Visium (10x Genomics) spatial transcriptomics slides (6 x 6 mm) in a cryostat and then fixed in methanol for 30 mins at -20 ºC. Tissue was incubated sequentially in haematoxylin for 7 min, bluing buffer for 2 min, and eosin for 1 min. Histology of stained tissue sections were imaged using a Leica SP8 microscope (Wetzlar, Germany, objective: HC PL APO 10x/0.40 CS2). Following imaging, sections were permeabilised for 18 mins, according to prior optimisation, and libraries were prepared per the manufacturer’s protocol. Libraries were pooled and sequencing was performed in a single run across 4 lanes on a NovaSeq 6000 v1.5 chemistry with an estimated library size of >115,000 read pairs per spot. In total, we detected 48,154 spots (2,645-4,422 per slide) with 109,893 (67,462-165,669) mean UMIs per spot and 90% of these mapped confidently to the genome (Supplementary Data 13).

#### Pre-processing

Read pairs were processed and aligned to a reference genome using Space Ranger (10x Genomics, Version 2.1.0) using the ‘count’ pipeline (genome build GRCh38, Ensembl 98). Aligned reads were preprocessed using Scanpy. Individual sections were normalised for library length (normalise_total), log transformed (log1p), and then concatenated into a unified dataset. The union of the 3000 most highly variable genes from each section were used to train the STAligner^87^ tool on the points from all slides, with prior calculated spatial networks (Cal_Spatial_Net [rad=300]). Mitochondrial and haemoglobin genes were omitted from training the model due to their association with sample quality. On the concatenated dataset, spots containing <100 genes, <100 total counts and/or >85% mitochondrial genes were removed (all spots were retained until the spatial integration was completed to retain integrity of spatial graphs). Neighbours were calculated using the STAligner output and the mclust algorithm was then used to cluster the spatial domains (7 clusters, selected based on layers I-VI and white matter). Spatial domains were visually confirmed across all sections to validate that conserved laminar structure was retained across individuals. Cluster/layer markers were determined using rank_genes_group (method = Wilcoxon). We were unable to robustly delineate layer I from layer II due to the thinness of this domain in BA11 but detected robust annotations for layers II-VI and white matter.

#### Single nucleus cell type spatial deconvolution

To map cell types from the whole snRNAseq dataset onto the spatial domains, the cluster mode of Tangram^88^ was used to predict the probability of cell types at each point of the integrated spatial dataset, using the intersecting 10,000 most highly variable genes over 1000 epochs. A benchmarking study has recommended Tangram for datasets where snRNAseq and spatial transcriptomics have been derived from the same individuals^23^. The calculated probability of each cell type at each spot was then multiplied by the total number of cells detected for that given cell type to obtain spatial and proportional distribution of cell types across the layers, relative to their abundance within the whole cell population.

To spatially resolve astrocyte subtypes, Tangram was used to impute the most probable layer onto each astrocyte nucleus. To prioritise astrocyte-relevant genes, genes with < 500 counts in < 1000 astrocyte nuclei were omitted before the clusters mode was trained on the intersecting 10,000 most highly variable genes from the spatial transcriptomics data over 1000 epochs, providing a probability of each cell occurring at each laminar domain. Each cell was then assigned to the most probable layer and cells with <1.5 fold change probability from other clusters were omitted for downstream analysis.

To explore the layer-specific astrocyte markers, a pseudobulked counts approach was performed similar to as performed for the snRNAseq dataset. No covariates were included in this model, given that each individual was equally represented between layer measurements.

### Immunohistochemistry

Fresh frozen OFC sections (10µm thick) were mounted onto SuperFrost microscope slides and were then thawed for 10 mins at 37ºC and post-fixed in 4% paraformaldehyde for 15 mins at room temperature (RT). Sections were then blocked using 10% Normal Donkey Serum (Merck), 1%w/v Bovine Serum Albumin (Sigma), and 0.3M glycine (Sigma) in TBS for 1h at 37ºC. Sections were incubated in primary antibody diluted to the corresponding ratio in antibody diluent: (1% BSA, 0.3% Triton X-100 in TBS) overnight at 4ºC (Supplementary Table 1). The following day, secondaries (Donkey anti-rabbit Alexa Fluor Plus 488 [A32790], Donkey anti-rabbit Alexa Fluor Plus 647 [A32787]) and HOECSHT336 (ThermoFisher) were diluted in antibody diluent and slides incubated for 1h at RT, treated with 0.03%w/v Sudan Black in 70% ethanol for 20 mins and coverslipped using Fluromount (Sigma). Each step was washed 3 x 5 mins in TBS-Tween 20.

#### Analysis of astrocyte morphology from immunohistochemistry

Tissue sections were imaged on a Leica DMi8 fluorescent microscope (Leica DFC9000 GTC camera: 4.2 MP sCMOS camera) with an LED source, fitted with THUNDER image clearing. Whole tilescan images of cortical tissue were cleared using ‘Large Volume Computational Clearing’. Astrocytes were detected using QuPath^89^ (version 0.4.4), controlling for detection parameters across samples (fluorescence threshold=500, minimum/maximum area=1000/200000µm^2^, sigma=6µm). Layer domains were identified using the morphological features of the nuclei and manually annotated. Cell density was measured as detected objects per annotation area. Area coverage of each identified cell was extracted per object and annotated per domain. For all statistical tests, linear regressions were performed. Variables were first tested for normality and non-normal data was transformed using a natural log transformation. Significant covariates were determined using a linear regression fitting all available full rank covariates, including Age, Sex, Postmortem Interval, RNA Integrity Number, and Brain pH. Covariates not significantly contributing to the model were omitted in the final fitted regression to increase the degrees of freedom. Multiple testing correction (Benjamini Hochburg, P_adj_ <0.05) was applied to all statistical tests. For the analysis of cell size, an inverse transformation was applied due to the strong right skew of the data and additional conservative multiple testing correction was applied, using a Bonferroni correction, with the number of cells being included in the number of tests (e.g. 6 comparisons * 1000 cells, n_tests=6,000). Individual was also included as a batch covariate for area measurements to account for biases in the contributions of cell numbers.

### Induced astrocyte differentiation for glucocorticoid activation

#### Stem cell maintenance

All experiments using human pluripotent stem cells (hPSCs) were approved by the University of Wollongong Human Research Ethics Committee (HREC 2020/451) and the Institutional Biosafety Committee (Exempt Dealing GT19/08). H9 hPSCs (WA-09, WiCell) were cultured in mTeSR1 media (Stem Cell Technologies, Vancouver, Canada) in feeder-free conditions on Matrigel (Corning, New York, United States) coated plates. Media was changed daily. When cells reached 70-80% confluency, culture was dissociated with Accutase (Life Technologies) and ∼10% of these cells were seeded as colonies. Thawed cells underwent two passages prior to the commencement of differentiation.

#### Lentiviral production

Lentivirus was produced using a 3^rd^ generation lentivirus system. Plasmid DNA (pVSVG [addgene #8454], pRSV-Rev [#12253], pMDL/pRRE [#12251] and either FUW-M2rtTA [#20342] or the SOX9/NFIB plasmid^44^ were complexed with PEI for 10 mins at 4^°^C at a ratio of 1:2.5. HEK293T cells greater than 80% confluency were transfected with plasmid complex for 2 h at 37^°^C and lentiviral particles collected as in previous description^90^.

#### Astrocyte differentiation

To generate induced astrocytes (iAsts), we utilised a system that exploits the over expression of transcription factors key for astrocyte lineage determination, as described in prior papers^44,91,92^. First, neural progenitor cells (NPCs) were produced according to previously described protocols^93,94^. Briefly, colonies of hPSCs were maintained until colonies were at least 70% confluent. To induce neural fate, media was switched to a DMEM base containing 100nM LDN193189 and 10µM SB431542, supplemented with 1X B27 and 1X Glutamax. After seven days, the media was switched to a media containing 20ng/mL EGF and 20ng/mL FGF2 for an additional seven days. The resulting NPCs were dissociated using Accutase and cryobanked.

To generate iAsts, NPC stocks were plated onto Matrigel-coated places in the EGF/FGF2 media alongside 1µM Y-27632 and both TTA and SOX9-NFIB lentiviruses (Day 14). The virus was removed after 12-16 h and media was replaced with Astrocyte Differentiation Media (1% B27 supplement, 2mM GlutaMAX, 1% FBS, 8ng/mL FGF-2, 5ng/mL CNTF, 10ng/mL BMP4) containing 2µg/mL Doxycycline (Day 15). To select for transduced NPCs, media was further supplemented with 2µg/mL puromycin (Day 16-19). On day 20, induced iAst lineage cells were replated for downstream applications in Astrocyte Differentiation Media supplemented with Y-27632. After 8-12h, media was replaced with Maturation Media (1% N2, 1mM Sodium Pyruvate, 2mM GlutaMAX, N-acetyl-cysteine 5µg/mL, 5ng/mL heparin-binding EGF-like factor, 10ng/mL CNTF, 10ng/mL BMP4, 100µM DBcAMP). iAsts were cultured in Maturation Media for an additional 15 days (days 20-35).

#### Dexamethasone stimulation

To model the glucocorticoid-mediated stress response in iAsts, glucocorticoid receptors were activated with 100nM equivalent dexamethasone (water-soluble) (Sigma-Aldrich, D2915) supplemented directly into Maturation Media. Cells were exposed for 0, 0.5, 1, 3, 5, or 7 days prior to endpoint (day 35) with complete media replacement every 48 h. All treatments had the same endpoint for downstream assays.

### Viability

Cell viability following dexamethasone treatment was determined using PrestoBlue (Invitrogen). Cells were incubated using a 1:10 dilution of reagent in Maturation Media for 30 mins at 37ºC. Viability was measured via fluorescence at excitation/emission of 560/590nm and normalised to wells containing only media.

### Gene expression (qPCR)

RNA isolation was performed using a modified TRIsure (Bioline) method. iAsts were washed once with PBS and then incubated in 500µL TRIsure per well of a six-well plate for five mins at RT and manually agitated using a cell scraper. Chloroform was added in a 1:5 ratio to cell lysate, mixed for 15 s and incubated for a further 15 mins at RT. Samples were then centrifuged at 12,000g for 15 mins at 4°C. The aqueous phase was removed, mixed 1:1 with isopropanol, and frozen overnight at -80°C to precipitate the RNA. The samples were thawed on ice and centrifuged at 12,000g for 10 mins at 4°C to pellet RNA. Pellets were washed with 900 µL 75% ethanol, centrifuged, and left to air dry. Dried RNA was resuspended in 20-50µL RNase-free water and further treated with a Turbo DNA-free kit as per instructions to remove genomic DNA. RNA was then treated with 10% 4M Sodium Acetate and 1:10 isopropanol and precipitated overnight at -80°C. Thawed RNA was pelleted at >20,000g for 20 mins at 4^°^C. Pellets were washed in 70% ethanol and centrifuged for a further two mins at >20,000g at 4°C. RNA was left to dry and resuspended in 20-50 µL RNase-free water. Yield and purity were determined using a NanoDrop 2000 spectrophotometer. One microgram of total RNA was reverse-transcribed to cDNA using the High-Capacity RNA-to-cDNA Kit (Thermo Fisher) and stored at -20^°^C. Quantitative PCR (qPCR) was performed using KiCqStart SYBR Green Master Mix (Merck) SensiFAST No-ROX Kit (Bioline). Additional RT-qPCRs were performed using pre-designed TaqMan assays with TaqMan Universal PCR Master Mix. Amplification was visualized and analysed using a Quantstudio5 system (Thermofisher). Gene expression was compared to the geometric mean the Ct of B2M, GAPDH, and HPRT1. All samples were run in triplicate for each probe.

### Immunocytochemistry

For all immunocytochemistry, cells were plated into optical glass bottom black-walled 96 well plates. At indicated timepoints, cells were washed three times with TBS and fixed with 4% PFA for 20 mins at RT, followed by another three washes in TBS. Cells were permeabilised for 5 mins in 0.5% Triton X-100. Samples were blocked for 1h in TBS containing 5% normal donkey serum and 3% BSA, and then incubated in primary antibody solution (as above for IHC) at the indicated dilutions overnight at 4^°^C (Supplementary Table 1). The following day, slides were washed three times with PBS and incubated in the respective secondary antibodies diluted 1:500 in antibody solution. Slides were washed three times and stored in a mounting medium consisting of 0.5% n-propyl gallate, 20nM Tris pH8.0, and 90% glycerol in water.

Cells were imaged using the same DMI8 setup as for the tissue but using small volume computational clearing. Positive cells were identified using QuPath, detecting the nuclei and expanding the radius 10µm and to capture the dual labelling in the cell body. Baseline settings were calibrated to control cells and maintained across all timepoints per protein visualised.

### Glutamate uptake assay

iAsts were plated into Matrigel-coated 48 well plates. On Day 20, media was removed, washed one with Hanks Buffered Salt Solution (HBSS) (140mM NaCl, 5mM KCl, 1mM CaCl, 0.5mM MgCl, 0.3mM NaPO4, 0.4mM KHPO4, 6mM Glucose, 4mM NaHCO3) and equilibrated in fresh HBSS for 20 mins at 37ºC. Media was then replaced with HBSS supplemented with 150uM L-Glutamic Acid (Sigma). Media was collected off the cells at 120 mins post-exposure and stored at - 20ºC. Controls consisting of Na+-free HBSS (NaCl replaced with equimolar Choline Chloride) and dlthreo-b-hydroxyaspartic acid, a potent inhibitor of EAAT family proteins, including EAAT1/2 were performed in parallel to validate efficacy of glutamate clearance. Cells were immediately lysed in ice cold Radioimmunoprecipitation Assay (RIPA) buffer and protein was quantified using a Bradford Assay (Bio-Rad), per manufacturer’s instructions.

Glutamate remaining in media was determined using a Glutamate Assay Kit (abcam, ab83389) following the provided protocol. Glutamate was subtracted from the glutamate treatment well containing no cells and normalized to the total protein of each well to determine relative glutamate uptake.

### Calcium imaging

Astrocytes seeded at 10,000 cells per well of a black-walled clear plastic bottom 96 well plate were washed and loaded with 4µM Fura-2, AM and 0.02% Pluronic Acid in HBSS for 45 mins at 37ºC. Cells were then washed for 10 mins in HBSS and placed in a starting volume of 80µL HBSS. Calcium activity was measured using a Flexstation 3. Respective compounds (20µL) [final conc] (ATP [30µM], glutamate [100µM], GABA [100µM], AMPA [100µM], NMDA [100µM]) were added at 20 seconds and ionomycin [3µM] was added after 4 mins followed by another 1 min of imaging. The F_340/380_ nm excitation ratio was measured every 4s at 510nm. Data was normalised to the first three readings (prior to compound addition). Peak response was taken as the highest normalised ratio measurement between 20 and 160 seconds. Maximum response was taken as the highest F_340/380_ ratio after the addition of ionomycin. To determine the proportion of the maximum response, normalised baseline was subtracted from all values and peak response was normalised to maximum response.

### Organoid-derived astrocytes

To validate cell findings, No.1 hiPSCs were reprogrammed from NuFF3-RQ newborn foreskin feeder fibroblasts of a male donor (GSC-3404, GlobalStem) and used to produce organoids as described previously^95^. Astrocytes were extracted from three separate batches of cerebral organoids. Cerebral organoids that were 375 to 446 days old were dissociated with a p1000 pipette and the cells were plated in Astrocyte Differentiation Media (1% B27 supplement, 2mM GlutaMAX, 1% FBS) supplemented with 10ng/ml EGF and 10ng/ml FGF-2. Cerebral organoids-derived cells were cultured for 2-3 days in the cerebral organoid medium (NDM+A medium: DMEM/F12GlutaMAX and Neurobasal in ratio 1:1, supplemented with 1:100 N2 supplement, 1:100 B27 with vitamin A, 0.5% nonessential amino acids, insulin 2.5 µg/mL, 1:100 antibiotic-antimycotic, and 50 µM 2-mercaptoethanol) and then in Astro-medium (in DMEM + Glutamax + 10% FBS + 1x Anti-Anti). The resulting astrocyte cultures (oAsts) were passaged using trypsin. oAsts were cultured until p3-p5. oAsts expressed the astrocyte markers SOX9, S100B and NFIA while they were negative for the neuronal marker MAP2. oAsts were exposed to either 100nM dexamethasone or vehicle (equal volume of DMSO) for three days. The amount of dexamethasone and timing was chosen as previously described^95^.

### RNA extraction

RNA and DNA were extracted from 18 samples (three technical replicates from three different batches of organoids, control and de×100nM groups) using the AllPrep DNA/RNA Micro kit from Qiagen (Cat. No 80284), nucleic acid concentration and quality were assessed with an Epoch plate reader. The mean 260/280 ratio for the RNA of all samples was 2.093 (minimum 1.831 and maximum 2.270). RNA sequencing library preparation was done with the NEBNext Ultra II Directional Library Prep kit for Illumina (Cat. No #E7765/L 96 reactions) using the NEBNext Poly(A) mRNA Magnetic Isolation Module (Cat. No E7490). Libraries were pooled according to their individual concentration and the pooling was assessed and re-calibrated with a test run in a MiSeq. The pool was sequenced in 3 lanes of the HiSeq4000 with paired-end 75bp run type and we acquired an average of 13,5 million reads per library. The samples were randomized in the library preparation plate taking into consideration their cell type origin, treatment group, RNA quality and cell culturing user to reduce as much as possible any technical variability.

### RNA sequencing

Differential expression analysis was performed using DESeq2^96^ with the standard settings. Genes related to glutamate regulation were extracted and analysed and GSEA was performed as described above.

## Statistical analysis

All statistical analysis was performed using R (version 4.2.1) using the RStudio environment (version 2023.12.1). Graphs were prepared either in Python (version 3.10.0) or R, dependent on the specific packages used. Cell data was presented as mean ± standard deviation unless otherwise stated. Statistical comparisons were performed using either linear regressions for two-group comparisons or one-way ANOVA paired with posthoc TukeyHSD testing for > two groups. All cell experiments were performed on 3 biological replicates. Further details of statistics are provided in respective figure legends and methods.

## Acknowledgements

We thank Dr. Simon Maksour and Dr. Neville Ng for optimising the protocol to derive iAsts, Dr. Amy Hulme and Dr. Tracey Berg for helping with plasmid design for the lentiviral delivery system, and Dr. Juan Pablo-Lopez for his critical feedback on the project and manuscript. We would also like to thank the donors for consenting to donate their brains to research, as well as the staff at the NSW Brain Tissue Resource Centre for their assistance with accessing this tissue. The NSW Brain Tissue Resource Centre is supported by the National Institute of Alcohol Abuse and Alcoholism of the National Institutes of Health under Award Number R28AA012725 and the Faculty of Medicine and Health, University of Sydney.

## Data availability

Key layer markers, as well as significant results from differential expression and gene set enrichment analysis are provided in Supplementary Tables. Pre-processed single-nuclei data is available at GEO (accession number: GSE254569). Spatial transcriptomics data will be made publicly available prior to publication. The remaining data is available upon request.

## Code availability

All code used to process, analyse, and display datasets, is available upon request.

