## Supplementary Figures for "Astrocytic glutamate regulation is shaped by adversity and glucocorticoid signalling"

a

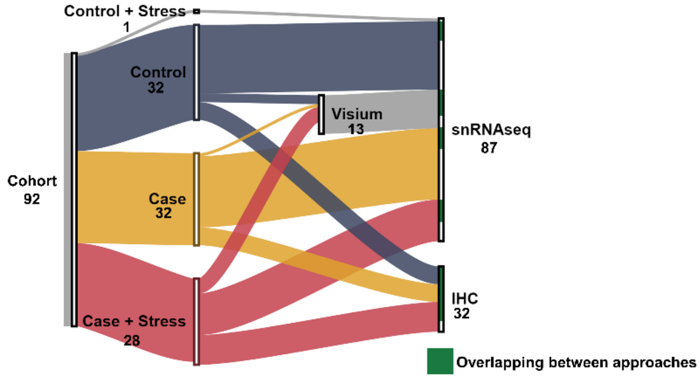


b

| Total *N* (snRNAseq/ IHC/ Visium) | Classification | Profound Psychological Stress | Diagnosis (SCZ/MDD/ BPD/SZA) | Age (years) (Mean±SD) | Sex (M/F) | PMI (h) (Mean±SD) | RIN (Mean±SD) | pH (Mean±SD) |
|---|---|---|---|---|---|---|---|---|
| **32**<br>31<br>8<br>3 | Healthy Control | No | - | **54.6±14.0**<br>57.3±12.7<br>49.0±17.9<br>55.1±15.7 | **20/12**<br>20/11<br>5/3<br>3/0 | **32.0±11.4**<br>30.6±11.0<br>34.4±4.4<br>32.2±10.2 | **7.2±1.3**<br>7.2±1.4<br>7.9±0.2<br>8.0±0.6 | **6.7±0.2**<br>6.7±0.2<br>6.7±0.1<br>6.8±0.1 |
| 1 | | Yes | - | - | - | - | - | - |
| **32**<br>32<br>8<br>2 | Psychiatric Disorder | No | 24/3/2/5 | **52.4±11.6**<br>52.8±11.2<br>60.0±0.0<br>50.8±11.1 | **19/13**<br>19/12<br>5/3<br>1/1 | **32.3±13.4**<br>31.8±13.7<br>30.5±4.2<br>31.6±9.7 | **7.3±1.5**<br>7.4±1.6<br>7.0±0.4<br>7.4±0.8 | **6.6±0.2**<br>6.6±0.2<br>6.5±0.1<br>6.5±0.1 |
| **27**<br>24<br>16<br>7 | | Yes | 17/5/3/3 | **52.5±16.4**<br>54.3±16.7<br>51.4±17.0<br>53.8±16.9 | **19/8**<br>16/8<br>12/4<br>7/0 | **37.3±19.0**<br>38.3±19.7<br>45.8±23.5<br>37.8±18.9 | **7.2±1.5**<br>7.4±1.3<br>7.4±1.3<br>6.8±1.7 | **6.6±0.3**<br>6.6±0.2<br>6.6±0.1<br>6.5±0.3 |

**Supplementary Figure 1: Demographics of cohort used in study. (a)** Alluvial plot illustrating breakdown of composition used for experiments. Green in final bar indicates proportion of individuals common to transcriptomic approaches and immunohistochemistry**. (b)** Table outlining key clinical covariates accounted for in this study. Abbreviations: SCZ: schizophrenia, MDD: major depressive disorder, BPD: bipolar disorder, SCA: schizoaffective disorder, M: Male, F: Female, PMI: postmortem interval, RIN: RNA integrity number.

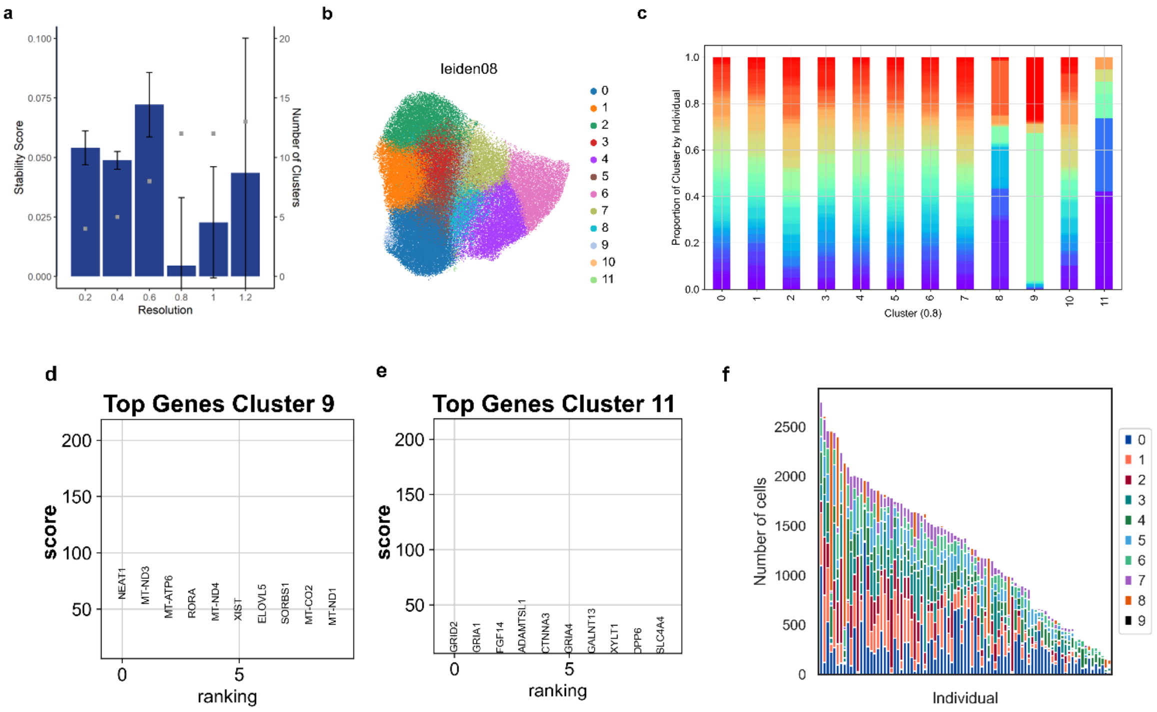


**Supplementary Figure 2: Extended profiling of snRNAseq astrocyte clustering. (a)** Cluster stability analysis using Leiden clustering at 0.2 increments. Deviation values between 20 iterations using the whole dataset and bootstrapping method run over 20 iterations using randomly selected n cells, where n cells are the proportion needed to remove a whole cluster (1 – 1/n clusters). Low value indicates better stability. Bars represent mean cluster stability (± 1SD). Grey points indicate number of clusters identified at each resolution. **(b)** Representative UMAP of initial clustering at selected resolution **(c)** Number of astrocytes per individual coloured by cluster number. **(d-e)** Top marker genes of removed clusters. **(f)** Cluster distribution per individual for tidied clusters.

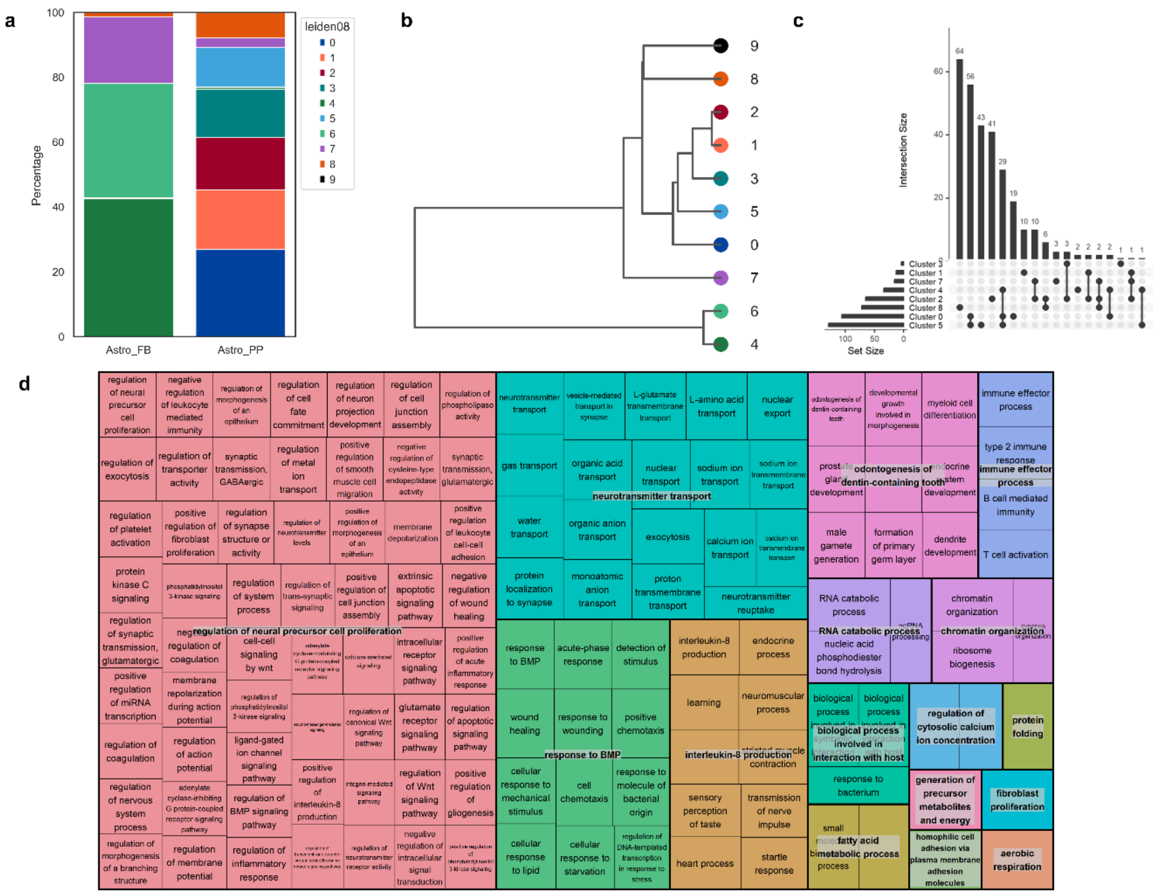


**Supplementary Figure 3: Extended profiling of astrocyte cluster identity. (a)** Proportion of astrocytes belonging to established identities. Broad cell types (Astro_FB/Astro_PP) were identified using label transferring approach on the whole cell population. Percentage of cells in each cluster belonging to broad identities as a percentage of all broad astrocyte cells plotted. **(b)** Dendrogram of cluster similarity based on the first 50 principal components. Pearson's correlation. **(f)** UpSet plot of intersection of C5 GO terms arising from gene set enrichment analysis. Set size indicates terms where $P_{adj} < 0.05$. Overlapping terms indicated as levels connected by a line and intersection size indicates terms associated with this intersection. **(d)** Semantic similarity of snRNAseq significant GSEA terms. ReviGO was used to cluster significant C5 gene ontology terms using the SimRel similarity measure. Terms with grey background denote family terms associated with terms sharing colour/thick border.

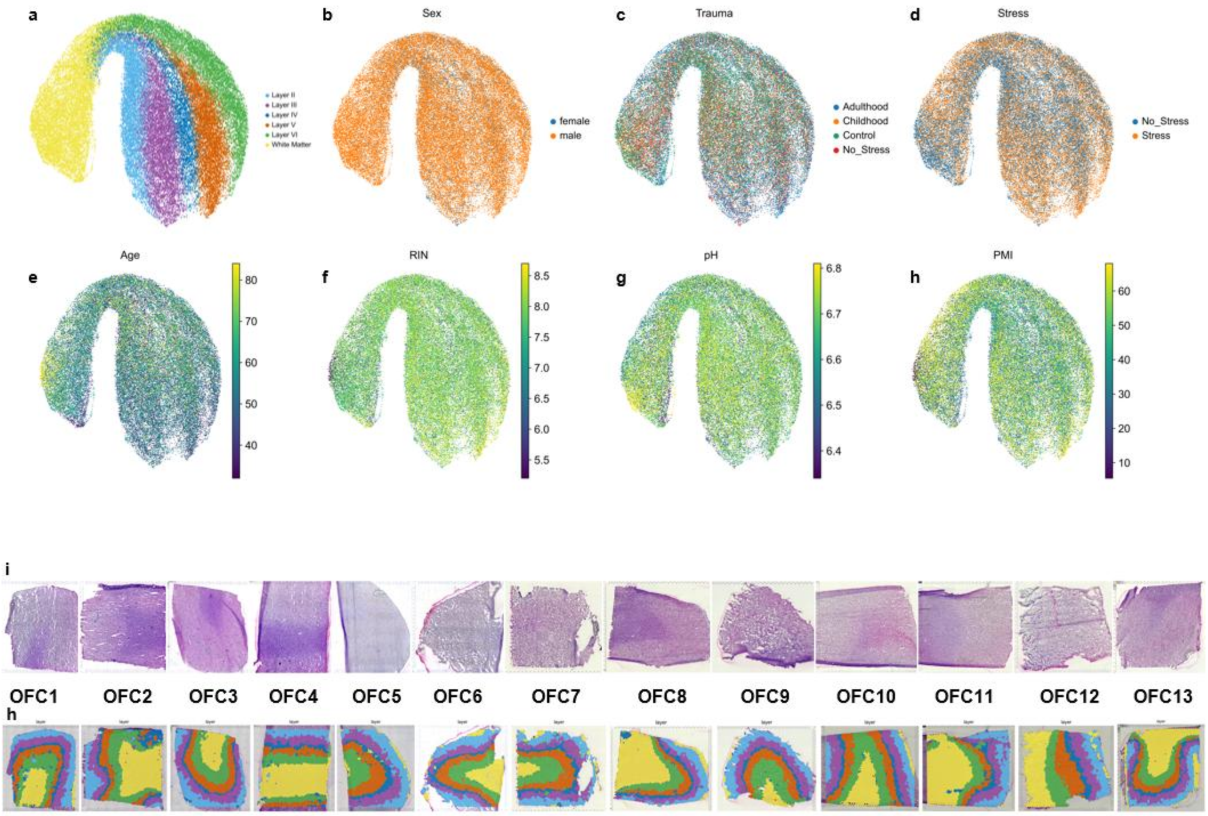


**Supplementary Figure 4: Spatial transcriptomics clustering validation. (a-h)** UMAP plot coloured by demographic covariates.**(i-h)** Histological validation of unbiased layer annotation. i) H+E staining of the capture regions containing the cortical sections used for spatial transcriptomics, demonstrating physical tissue quality across the cohort. h) remapping the unbiased integration of layer annotations back on to all sections used within the cohort demonstrates consistent laminar identification across the orbitofrontal cortex.

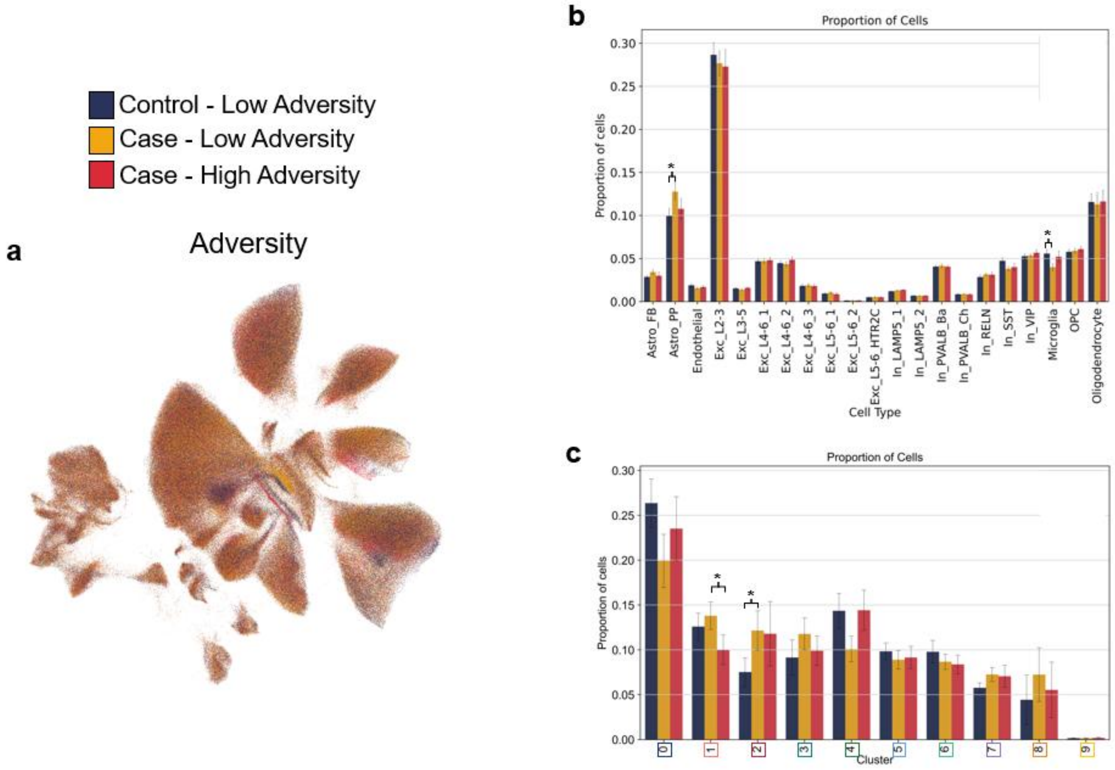


**Supplementary Figure 5: Single nucleus RNAseq cell type proportions delineated by adversity stratification. (a)** 2D UMAP representation of snRNAseq dataset coloured by adversity stratification. No cell type demonstrates biased clustering from any group. **(b)** Proportion of broad cell types grouped by adversity stratification. Significance calculated using a linear regression correcting for sequencing batch. **(c)** Proportion of astrocyte subclusters grouped by adversity stratification. Coloured border indicates cluster. Significance calculated using a linear regression correcting for sequencing batch. *$P_{nom}$<0.05.

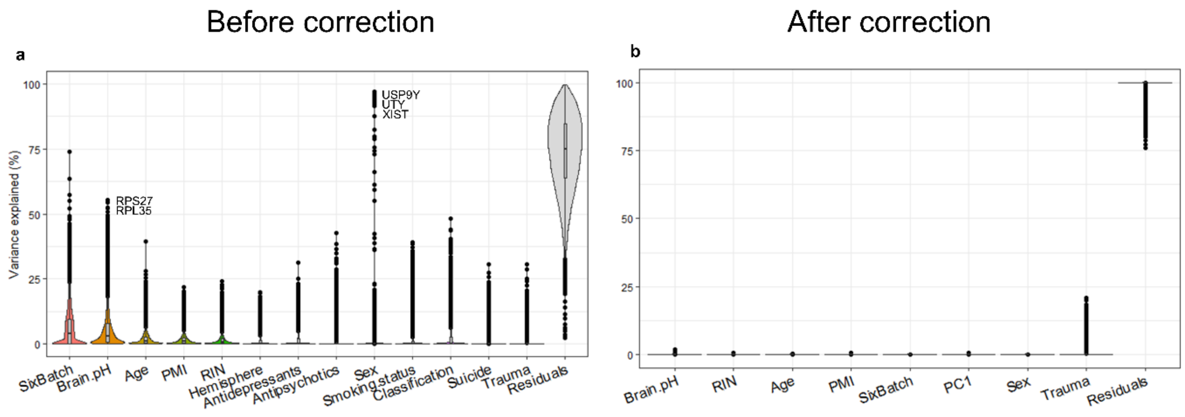


**Supplementary Figure 6: Variance partitioning of single nucleus RNAseq. (a)** Variance partitioning using pseudobulk counts for all astrocytes before batch correction. Representative genes demonstrate faithful partitioning of covariate-associated genes. **(b)** Variance partitioning using pseudobulk counts for all astrocytes after batch correction demonstrates that the method reduces both covariate noise as well as unexplained residuals, corrected for with PC1.

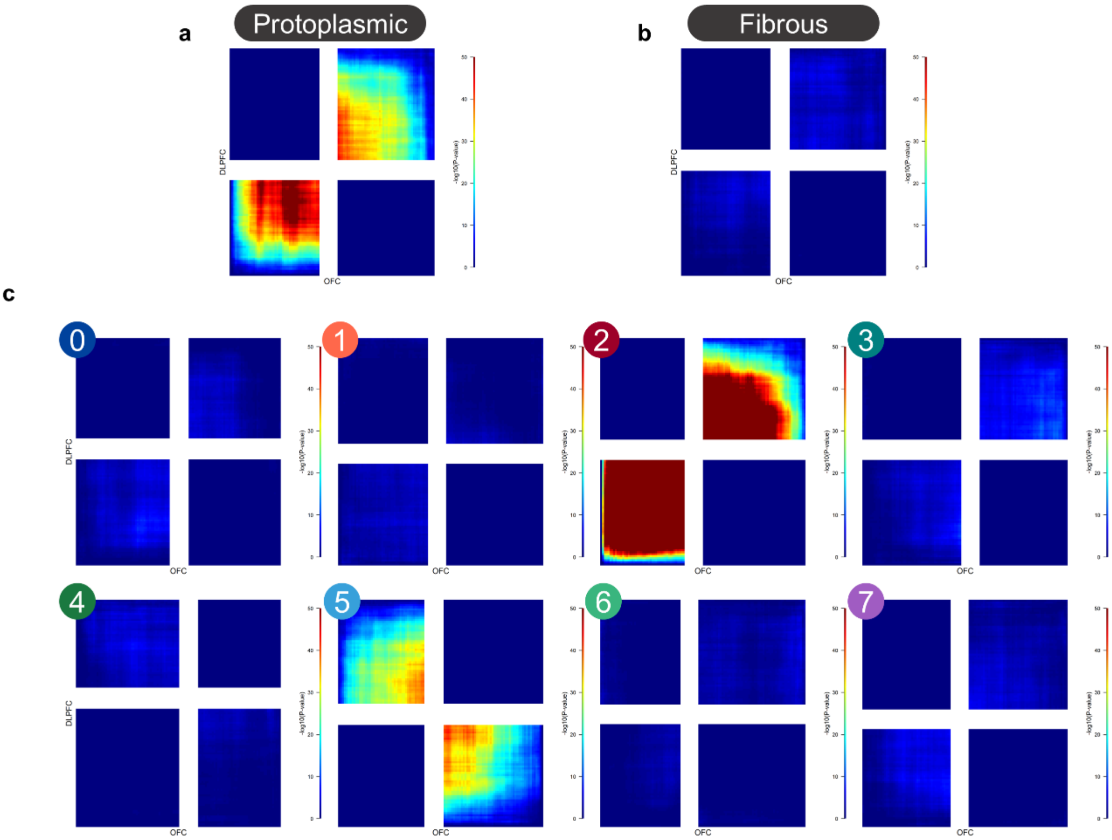


**Supplementary Figure 7: Rank rank hypergeometric overlap between DEGs from primary cohort (x-axis) to validation cohort (DLPFC, y-axis) (high-adversity-cases vs low-adversity-cases) by cluster. a)** Label transferring from the primary cohort to the replication cohort for protoplasmic astrocytes (Allan Brain Atlas) **b)** Label transferring from the primary cohort to the replication cohort for fibrous astrocytes (Allan Brain Atlas) **c)** Label transferring from the primary cohort to the replication cohort for defined astrocyte clusters. Scale set to -log10P 0-50.

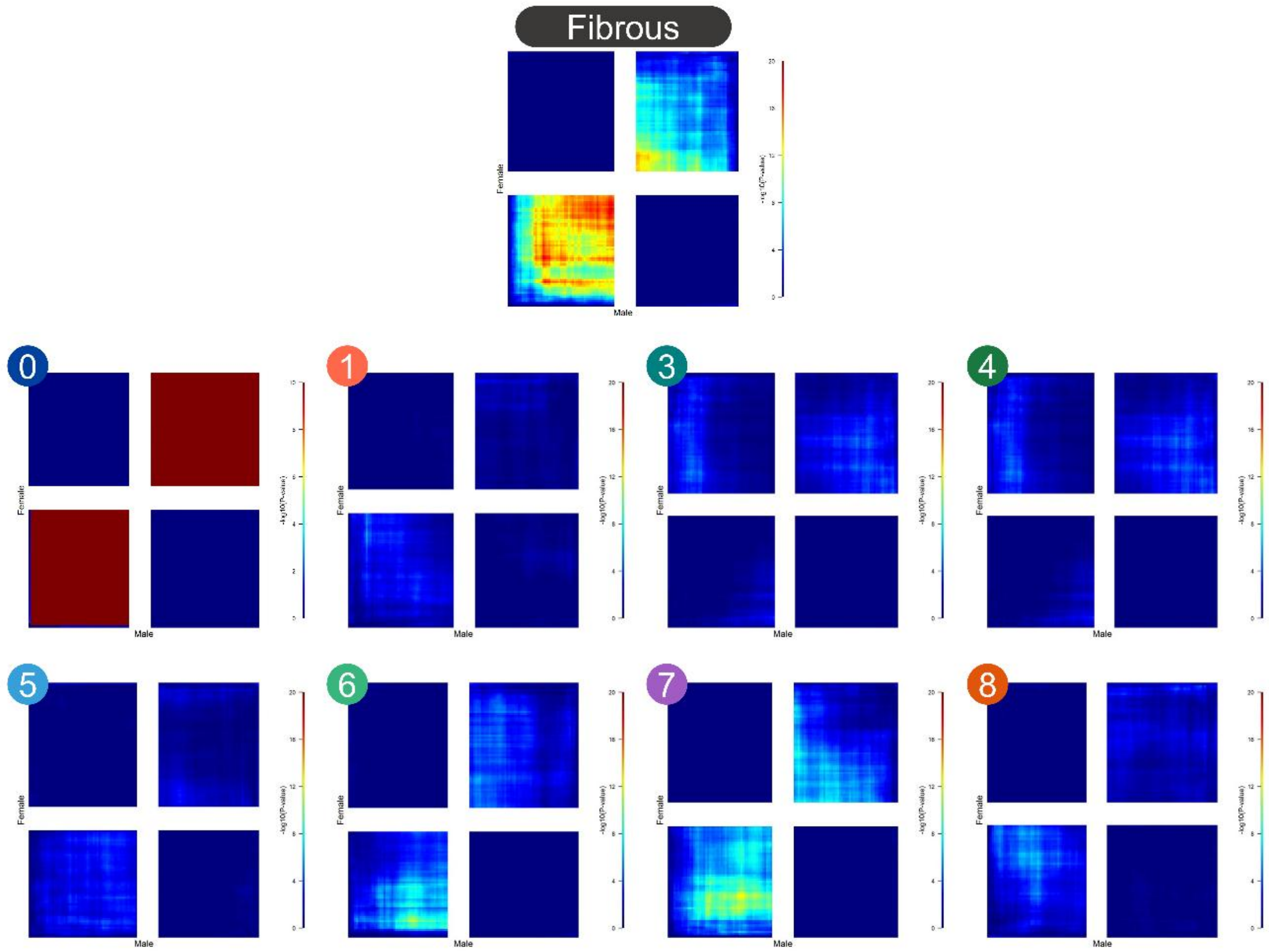


**Supplementary Figure 8: Rank rank hypergeometric overlap between male and female DEGs (high-adversity-cases vs no low-adversity-cases) by cluster.** Scale set to -log10P 0-20.

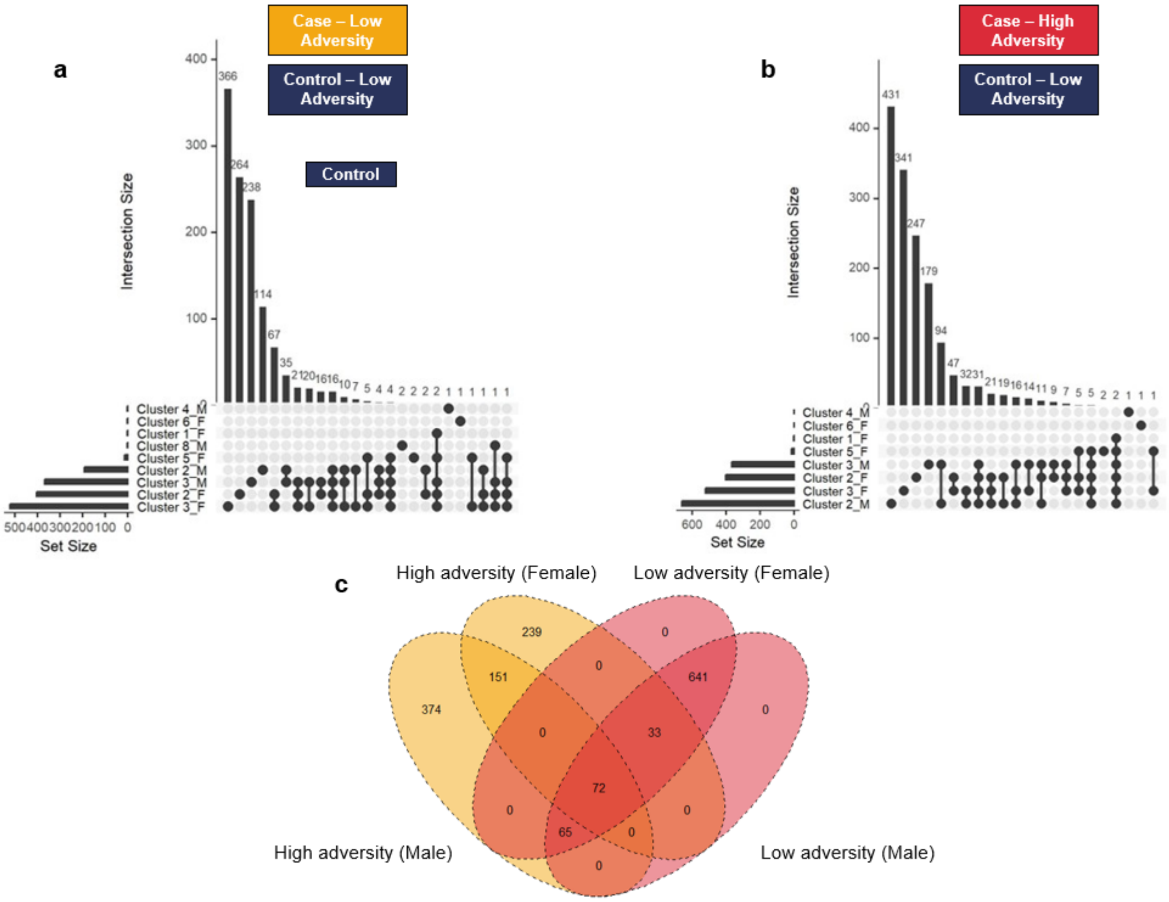


**Supplementary Figure 9: Differentially expressed genes between cases and controls by sex. (a)** UpSet plot of DEGs between low-adversity-cases and low-adversity-controls by sex ($P_{adj}<0.1$). Set size indicates number of genes meeting threshold for each cluster and dots connected by a line indicate common genes. **(b)** UpSet plot of DEGs between high-adversity-cases and low-adversity-controls by sex ($P_{adj}<0.1$). Set size indicates number of genes meeting threshold for each cluster and dots connected by a line indicate common genes. **(c)** Venn diagram of total DEG overlap by profound adversity (adversity) or not (case).

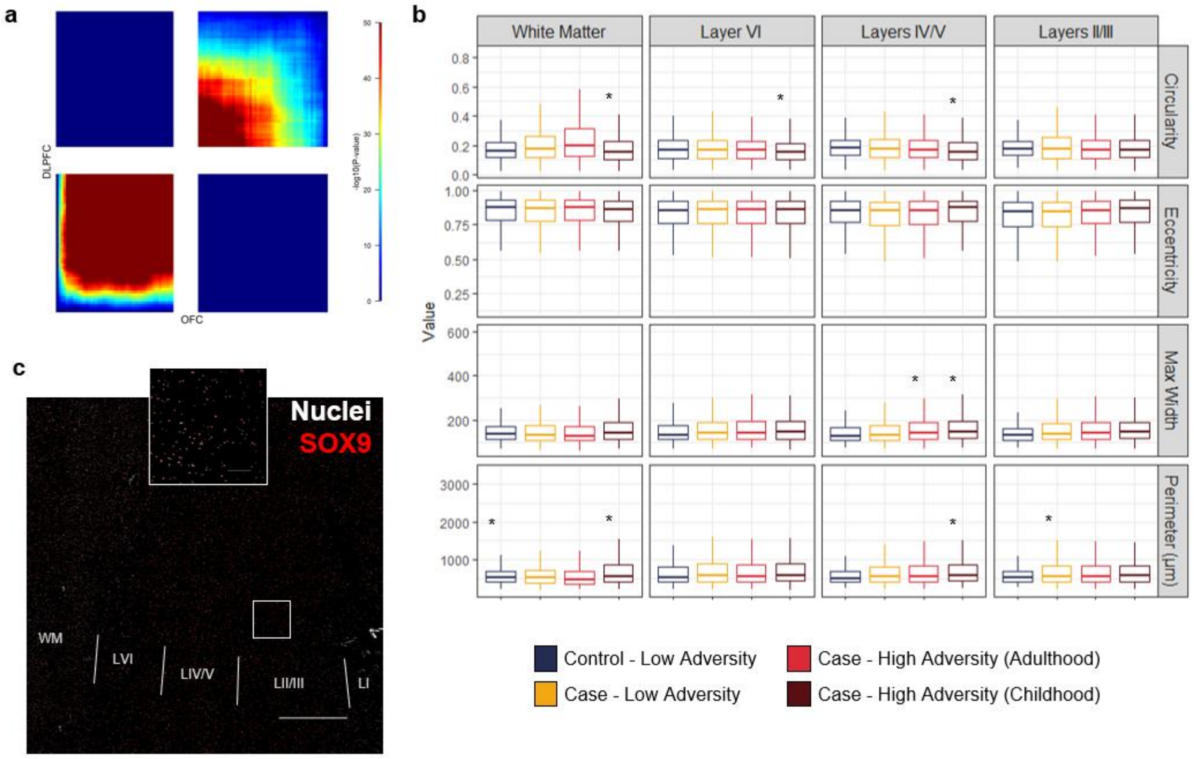


**Supplementary Figure 10: Supplementary Adversity Timing Testing a)** Rank rank hypergeometric overlap between adversity timing in cluster 2 in the primary cohort and the replication cohort. Scale = -log10(P) 0-50. **b)** Additional morphology measurements made. Significance indicates difference from low-adversity-cases. **(b)** Selected region demonstrating close-up staining. Scale bar on tilescan indicates 1mm, and 100µm in magnified region.

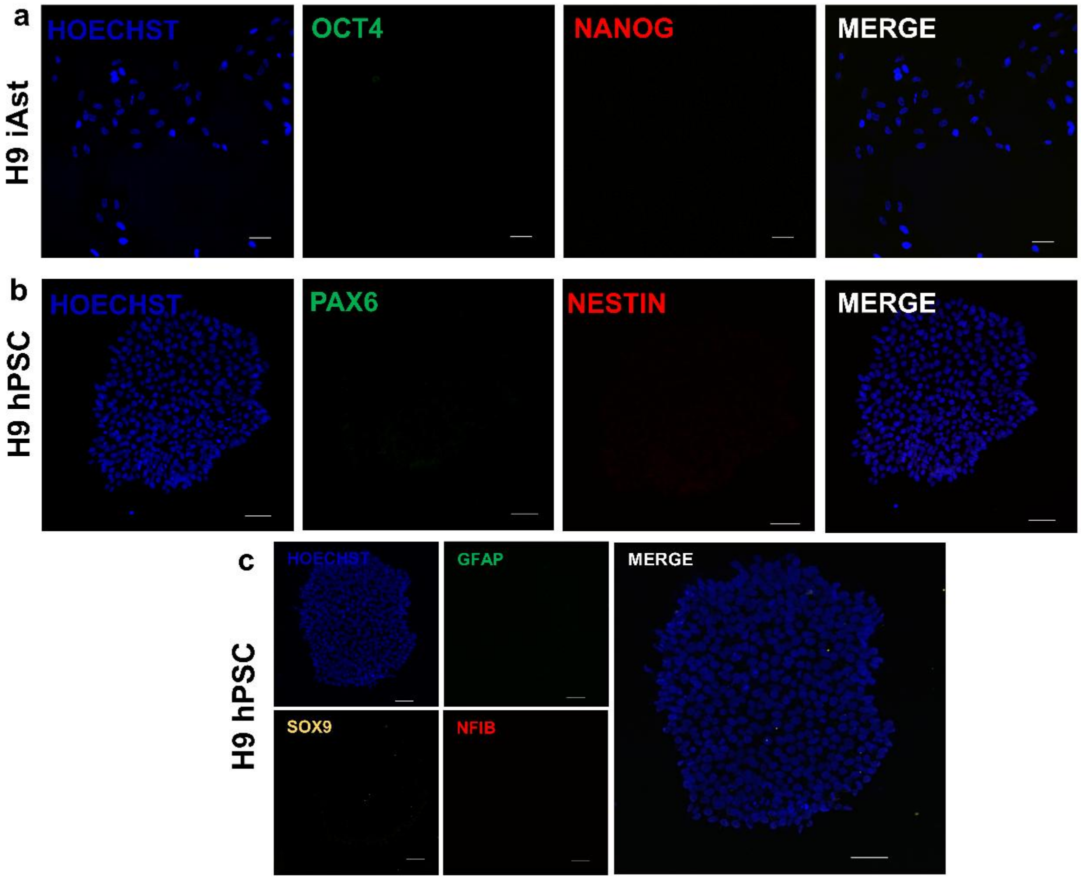


**Supplementary Figure 11: Immunocytochemical negative controls for cell types. (a)** Validation that iAst do not express pluripotency markers, OCT4 (green) and NANOG (red). **(b)** Validation that H9 human pluripotent stem cells do not express neural progenitor markers, PAX6 (green) and NESTIN (red). **(c)** Validation that H9 human pluripotent stem cells do not express astrocyte markers GFAP (green), SOX9 (yellow) and NIFB (red). Primary antibodies were used at the same concentration as outlined in Supplementary Table 1 and positive controls are available in Figure 6. Scale bar indicates 50µm.

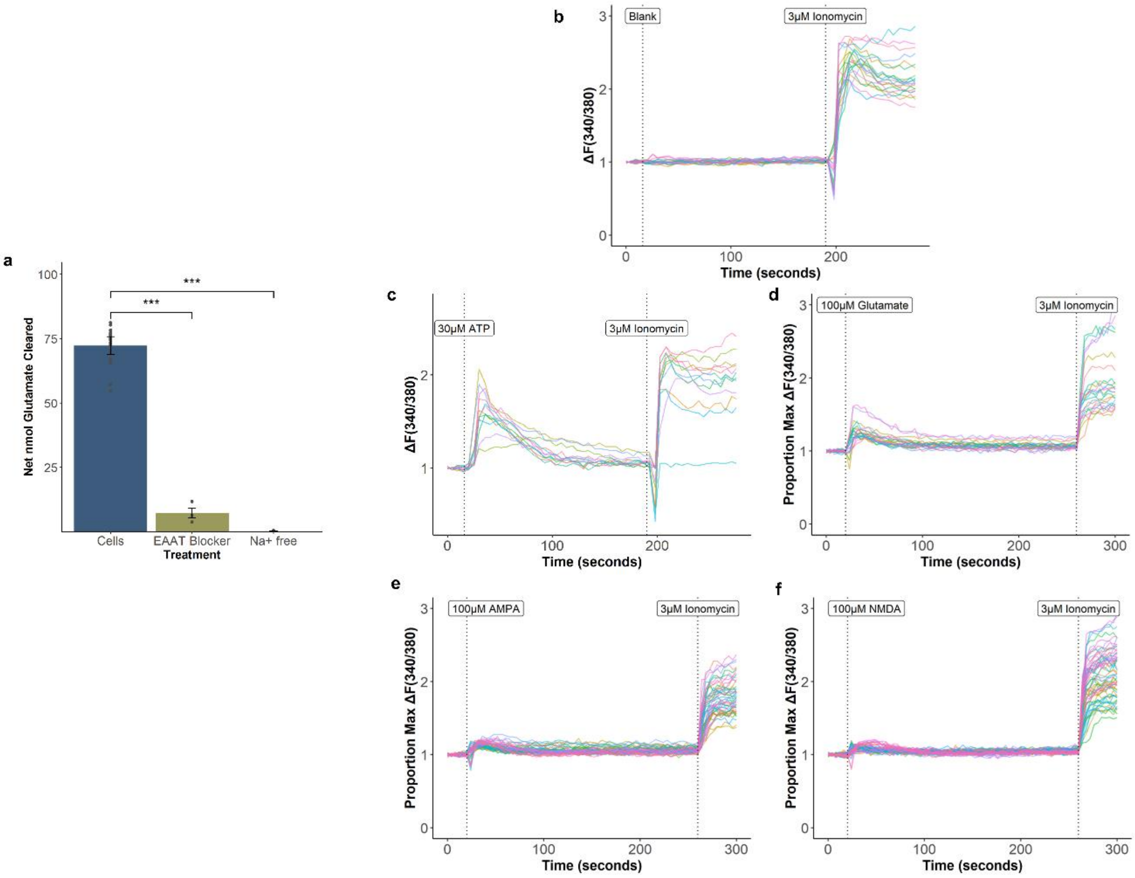


**Supplementary Figure 12: Further validation of cell model. (a)** Glutamate clearance of iAsts over 120-minute period. EAAT blocker DL-TBOA, NA$^+$ free, HBSS, NaCl replaced with equimolar choline chloride. Each point indicates an individual well experiment.DL-Bars indicate means ± 1 SD.***P<0.001 (ANOVA). **(b-f)** Flexstation time-response plots showing 340/380 emission ratio in response to various agonists. Plots were normalised to the mean of the first three recordings prior to addition of first agonist. Each line indicates the response of an individual well. N=3 biological replicates, each with two technical replicates.

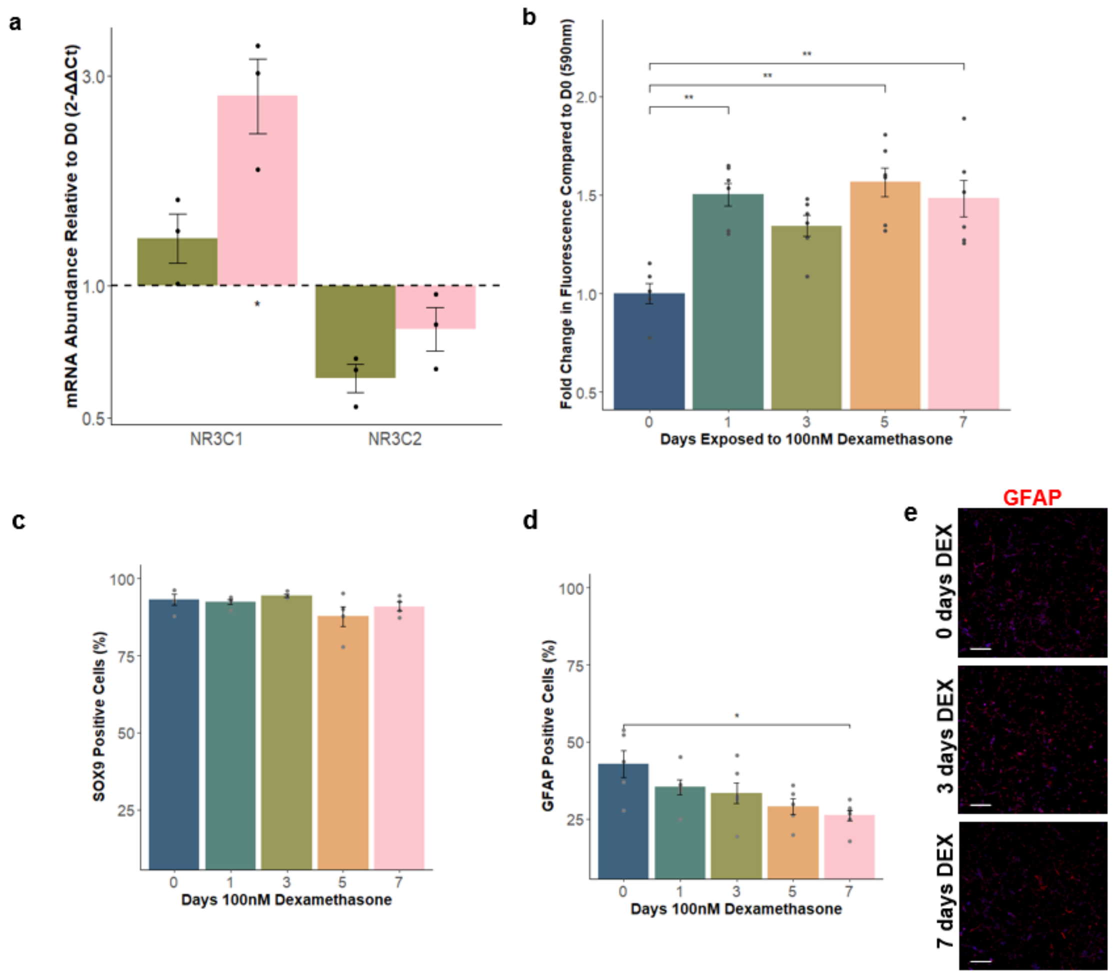


**Supplementary Figure 13: Additional validation of glucocorticoid-mediated biological paradigm. (a)** qPCR of glucocorticoid receptors in response to 100nM dexamethasone. Significance relative to vehicle. **(b)** Presto blue assay to measure cell viability. Bars indicate mean ± SD. Each point represents and individual well. N=3 biological replicates, each with two technical replicates. **c)** Quantification of SOX9+ astrocytes (as proportion of number of nuclei) over different timepoints of DEX exposure. **d)** Quantification of GFAP+ astrocytes (as proportion of number of nuclei) over different timepoints of DEX exposure. **e)** Representative staining of GFAP at different timepoints of exposure (Scale = 50µm). *P<0.05, **P<0.01 (ANOVA, TukeyHSD).

**Supplementary Table 1: Antibody information for antibodies used.** Abbreviations: ICC, immunocytochemistry; IHC, immunohistochemistry

| Antibody Target | Catalogue # | Company | Dilution | Species |
|---|---|---|---|---|
| EAAT2 | ab41621 | Abcam | IHC: 1:200<br>WB: 1:5000 | Rabbit |
| SOX9 | ab76997 | Abcam | IHC: 1:200<br>ICC: 1:500 | Mouse |
| AGT | AF3156 | R&D Systems | ICC: 1:200 | Goat |
| EAAT1 | ab416 | Abcam | ICC: 1:200 | Rabbit |
| GS | GTX109121 | GeneTex | ICC: 1:500 | Rabbit |
| GFAP | AB5541 | Merck Millipore | ICC: 1:1000 | Chicken |
| FKBP5 | sc-271547 | Santa Cruz | ICC: 1:100 | Mouse |
| GAT3 | ab431 | Abcam | ICC: 1:200 | Rabbit |
| ALDH1L1 | GTX84892 | GeneTex | ICC: 1:100 | Mouse |
| S100B | Ab52642 | Abcam | ICC: 1:100 | Rabbit |
| NESTIN | ABD69 | Merck Millipore | ICC: 1:500 | Rabbit |
| PAX6 | AB_528427 | DSHB | ICC:1:50 | Mouse |
| OCT4 | Ab181557 | Abcam | ICC: 1:200 | Mouse |
| NANOG | Ab21624 | Abcam | ICC: 1:1000 | Rabbit |
